# A novel family of lncRNAs relate facioscapulohumeral muscular dystrophy to nucleolar architecture and protein synthesis

**DOI:** 10.1101/2024.06.29.600824

**Authors:** Valentina Salsi, Francesca Losi, Bruno Fosso, Marco Ferrarini, Sara Pini, Marcello Manfredi, Gaetano Vattemi, Tiziana Mongini, Lorenzo Maggi, Graziano Pesole, Anthony K. Henras, Paul D. Kaufman, Brian McStay, Rossella Tupler

## Abstract

Facioscapulohumeral muscular dystrophy (FSHD) is a hereditary myopathy linked to deletions of the tandemly arrayed D4Z4 macrosatellite repeats at human chromosome 4q35. These deletions accompany local chromatin changes and the anomalous expression of nearby transcripts *FRG2A, DBET,* and *D4Z4.* We discovered that *FRG2A* is one member of a family of long non-coding RNAs (lncRNAs) expressed at elevated levels in skeletal muscle cells with distinct amounts detected in individual patients. We found that *FRG2A* lncRNA preferentially associates with rDNA sequences and centromeres and promotes the three-dimensional association of centromeres with the nucleolar periphery in FSHD cells. Furthermore, we demonstrate that the elevated *FRG2A* expression in cells from FSHD patients reduces rDNA transcription and global protein synthesis. Our results frame an entirely unanticipated new disease model in which elevated lncRNAs levels mediated by deletions of D4Z4 macrosatellite repeats leads to a diminished protein synthesis capacity, thereby contributing to muscle wasting.

## Main

Two percent of the genome is composed of tandemly arrayed repeats of large size (TAR_ls_) commonly known macrosatellites. The organization, number and arrangement of these repeats was impossible to interrogate until the advent of long-read sequencing and the assembly of a complete human genome from telomere to telomere (T2T) ^1^. Recent research suggests that TAR_ls_ play essential roles in the maintenance of genome homeostasis by altering genome architecture, epigenome markings, and the regulation of gene expression^2,3^. However, the relationships between TAR_ls_ copy number variations and human physiology and disease are still unclear^2^.

Although TAR_ls_ are not sequence-related, they share several common characteristics: they extend in tandem over hundreds of kilobases, encompassing significant portions of the genome; they are rich in CpG sites, making them often subject to regulation by DNA methylation; and they frequently express both noncoding and coding RNAs. Collectively, it is now recognized that TAR_ls_ play a structural and regulatory role in chromatin organization within the nucleus^2^.

Facioscapulohumeral muscular dystrophy (FSHD) (MIM 158900) pathogenesis is currently linked to deletions of complete copies of the tandemly arrayed D4Z4 macrosatellite ^4–7^. A pathogenic threshold number of D4Z4 repeats (≤ 10) has been established and used for FSHD diagnostics^6,7^. Importantly, large clinical datasets based on a standardized clinical classification ^8^ have been generated and these studies concluded that FSHD clinical presentation varies widely ^9–14^. A rough genotype-phenotype correlation between the number of repeats and the severity of the disease holds well among those who carry deleted alleles with 1 to 3 repeat units. However, this correlation is often broken by high levels of intrafamilial and interindividual clinical heterogeneity, especially in patients with intermediate-sized alleles^11^ (4-10 repeats). Furtehrmore, D4Z4 alleles with 10 or fewer repeats have been detected in healthy carriers (4.6%)^15^, and in individuals with other myopathies. Therefore, D4Z4 array deletions contribute to a complex mechanism that only in certain, still unknown, conditions, leads to muscle wasting.

The prevalent model is that D4Z4 deletion causes *in cis* epigenetic changes at the 4q subtelomere resulting in transcriptional derepression of proximal genes^16–19^. Consistent with this view, we recently reported that the 4q subtelomere is subdivided into discrete domains, each with characteristic chromatin features associated with distinct gene expression profiles^20^. These discrete domains undergo diverse region-specific chromatin changes upon treatment with chromatin enzyme inhibitors or genotoxic drugs. Upon DNA damage, the 4q35 telomere-proximal *FRG2 (*hereafter *FRG2A), DBE-T* and *D4Z4*-derived transcripts are induced to levels inversely correlated with the D4Z4 repeat number. All these transcripts are normally barely detectable but stabilized through post-transcriptional mechanisms and bound to chromatin^20^. Among these, *FRG2A* exhibits the strongest response to external stimuli, showing the higher variations in its levels in respect to the others 4q35 genes. Additionally, FRG2A transcript is abnormally elevated in FSHD muscle samples^17^ and increases during myoblast differentiation to myotubes^21^. Several studies also reported that *FRG2A* expression during myogenic differentiation is regulated by long-range chromosomal interactions involving the 4q35 subtelomere^22,23^. However, although *FRG2A* expression is regulated by environmental, developmental, and genetic stimuli, the biological features of *FRG2A* gene products are poorly understood, and no evidence has been provided regarding the function or localization of FRG2A-encoded proteins.

Here, we demonstrate that *FRG2A* functions as a long non-coding RNA. We describe an unforeseen role of *FRG2A* in the regulation of rRNA transcription in muscle cells, which in turn affects the global rate of protein synthesis. We also report the unexpected impairment of nucleolar function in FSHD, a feature that could be shared by other myopathies and involved in muscle wasting during ageing.

### The T2T-CHM13 human genome assembly contains fourteen FRG2-like sequences

As the first step to investigate the biological role of *FRG2A,* we interrogated the T2T-CHM13 human genome assembly and found 14 *FRG2* paralogs (*FRG2s*; Figure 1a), while only six were annotated in the previous GRCh38 assembly. In the T2T genome, we detected six complete copies (2918 bp from TSS to polyA signal), located on chromosomes 4 (*FRG2A*), 10 (*FRG2B*), 20 (*FRG2EP*) and 22 (*FRG2C1*) and in two different positions on chromosome 3 *(FRG2C* centromeric and *FRG2FP* telomeric). Out of the 14 *FRG2* paralogs, four have predicted protein coding potential *(FRG2A*, *FRG2B*, *FRG2C* and *FRG2C1)*, while the others are annotated as “unprocessed pseudogenes” based on CAT-Liftoff gene annotation^24,25^. Predicted proteins encoded by paralogs with coding potential display distinct amino acid sequences and lengths (284aa FRG2C; 281aa FRG2A and B, 164 aa FRG2C1) (Supplemental Figure 1).

**Fig. 1.**
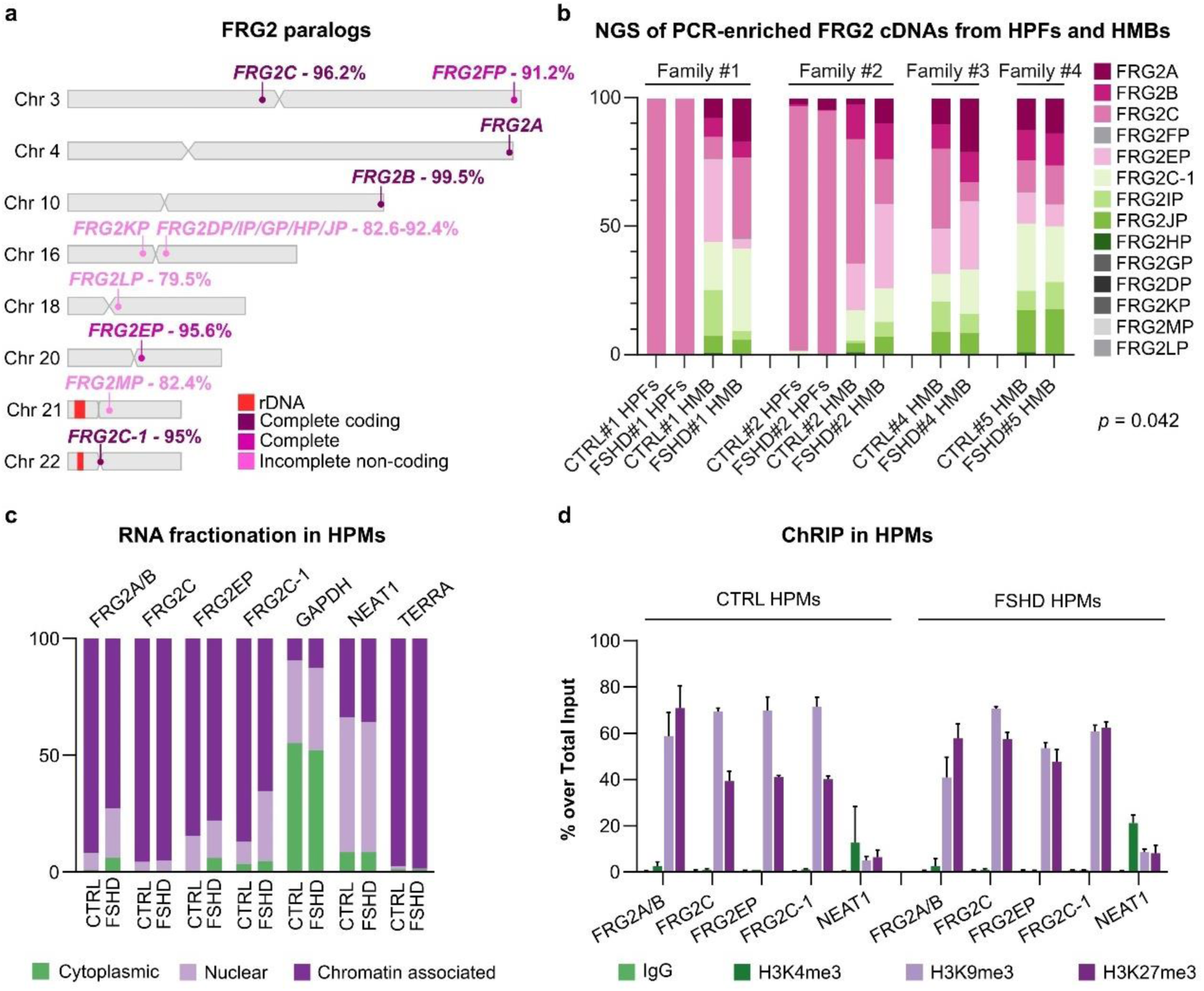
The human genome encodes fourteen FRG2 paralogs. **a**, Chromosome ideograms showing the location of *FRG2* paralogs and percentage identity to *FRG2A* on chromosome 4. The different colors indicate *FRG2* paralogs that are complete (2918bp) with coding potential (dark purple) (*FRG2A* ENSG00000205097.6, *FRG2B* ENSG00000225899.8, *FRG2C* ENSG00000172969.7, *FRG2C-1* ENSG00000172969.7) complete without coding potential (magenta) (*FRG2FP* ENSG00000232783.5, *FRG2EP* ENSG00000282842.1) or partial and non-coding (light pink) (*FRG2KP* ENSG00000261741.5, *FRG2DP* ENSG00000261711.6, *FRG2IP* ENSG00000260598.1, *FRG2GP* ENSG00000260291.1, *FRG2HP* ENSG00000260846.2, *FRG2JP* ENSG00000259897.2, *FRG2LP* ENSG00000267704.1, *FRG2MP* ENSG00000275170.1) according to T2T-CHM13 annotation. **b**, Genomic origin of FRG2s transcripts: NGS of PCR-enriched *FRG2* cDNAs obtained from human primary fibroblasts (HPFs) and human muscular biopsies (HMBs) from FSHD subjects and healthy relatives from 4 families. The analysis demonstrated muscle-specific expression of FRG2A and its overexpression in FHSD HMBs compared to controls. The statistical significance of FRG2A expression was assessed using a paired t-test, comparing the mean number of reads in controls and FHSD HMBs (*p* = 0.042). **c**, RNA fractionation experiment conducted in CTRL and FSHD-derived human primary myoblasts (HPMs). The predicted protein-coding FRG2 paralog transcripts were enriched in the chromatin-associated fraction of the RNA. GAPDH, NEAT1^101^ and TERRA^34^ were used as positive controls for cytoplasmic, nuclear and chromatin-associated RNA enrichment, respectively. **d**, Chromatin-RNA immunoprecipitation (ChRIP) in CTRL and FSHD HPMs. Chromatin was immunoprecipitated with the indicated antibodies. both in control and FSHD cells. Enrichment of target transcripts is shown as percentage over total input. FRG2s-t were enriched with heterochromatin-associated histone marks H3K9me3 and H3K27me3, NEAT1 was used as a negative control because it associates with H3K4me3-marked euchromatin^101^.

To determine which paralogs encode stable transcripts, we performed NGS sequencing of PCR-enriched *FRG2* cDNAs, comparing muscular biopsies as well as fibroblasts from healthy control individuals and FSHD patients from four families (Supplemental Table 1). Taking advantage of chromosome-specific SNPs, we identified the genomic loci encoding each FRG2 transcript (Figure 1b and Supplemental Figure 2). In the fibroblast samples *FRG2C* transcripts were detected almost exclusively. In contrast, transcripts from many *FRG2* paralogs were detected in the muscle samples, and their relative expression levels differed among individuals. This suggests a regulation mechanism that is both tissue-specific and subject-specific. Consistent with previous reports^20,21,26^, this analysis confirmed that *FRG2A* is expressed only in muscle cells and significantly overexpressed in FSHD-derived samples in comparison to healthy control family members (*p*=0.042, Figure 1b).

We observed that five out of the nine FRG2 paralogs that produce transcripts—namely FRG2FP, FRG2GP, FRG2HP, FRG2IP, and FRG2JP—lack predicted coding potential. Most of these genes are located in the pericentromeric region of chromosome 16 and comprise only incomplete sequences without an open reading frame (ORF).

Additionally, FRG2EP and FRG2FP, despite being transcribed, would encode very short proteins of just 10 and 79 amino acids, respectively, if translated. We thus concluded that *FRG2* paralogs can produce transcripts in primary cells and in muscle biopsies, some of them lacking coding potential.

### *FRG2* paralogs encode a novel family of heterochromatin-associated long non-coding RNAs

Having detected both coding and non-coding *FRG2* transcripts, we next tested whether we could detect *FRG2*-encoded proteins. To overcome the lack of available antibodies that recognize FRG2 proteins, we created 4 different edited cell lines with FLAG epitope tag sequences in frame with *FRG2* ORFs. Via CRISPR-Cas9 technology (Supplemental Figure 3A) we placed a FLAG epitope sequence at either the N-terminus or the C-terminus of *FRG2* genes, in both HeLa and HEK293 cells. In bulk populations, we observed the presence of Flag-tagged-knocked-in alleles at *FRG2A* (4q35), *FRG2B* (10q26), *FRG2C* (3p12), *FRG2C1* (22p11) coding-predicted genes and at *FRG2EP* (20q11) and *FRG2FP* (3q29) (Supplemental Figure 3b-e). Specific primers for each coding-predicted paralog were designed and tested using CHO-monohybrid cell lines that contain a single human chromosome (Supplemental Figure 3f). In all the edited cell lines, both the endogenous (Supplemental Figure 3g) and the Flag-tagged FRG2 transcripts (Supplemental Figure 3h) were easily detectable and, as expected, their expression increased significantly after doxorubicin treatment^20^ (Supplemental Figure 3i).

However, immunoblotting using the antibody against the Flag epitope failed to detect any FRG2 proteins (Supplemental Figure 4). The inability to detect FRG2 proteins by immunoblotting raised the question whether FRG2 transcripts are translated into proteins. We thus interrogated multiple databases for gaining evidence of FRG2 protein expression. First, we examined the Genome Wide Information on Protein Synthesis (GWIPS) database^27,28^ that cataloged translational potential via ribosome profiling^27^ and searched within deposited available RiboSeq data. The first database did not detect FRG2 RNA association with ribosomes (Supplemental Figure 5) or the presence of *FRG2*-derived peptides. We also failed to detect FRG2 peptides within available proteomic databases from human myoblasts^27,28^. Likewise, the Protein Atlas^29^ and Uniprot^30^ databases lack *FRG2*-derived proteins identified in proteome samples. We also performed an untargeted proteomic analysis in HeLa cells with cells treated with doxorubicin that induce stabilization of FRG2 RNAs^20^. More than 6,000 human proteins were identified, but FRG2s were not detected (Supplementary Table 2). Collectively, these data suggested that FRG2 transcripts might be rarely if ever translated into proteins.

We therefore focused on FRG2s-RNAs to learn about their potential as regulatory molecules. First, we performed RNA extraction upon cell fractionation experiments, in HeLa cells and primary biopsy-derived myoblasts from control or FSHD-affected individuals (Figure 1c and Supplemental Figure 6a). In both primary myoblasts and HeLa cells, transcripts from *FRG2* paralogs were highly enriched in the chromatin-associated fraction. To profile the types of chromatin interacting with FRG2s transcripts, we performed Chromatin-RNA Immunoprecipitation (ChRIP)^33^ experiments using antibodies raised against histone marks typical of euchromatin or heterochromatin (Figure 1d and Supplemental Figure 6b). We found that FRG2s transcripts are enriched in H3K9me3- and H3K27me3-marked but not H3K4me3-marked fractions, indicating that they are preferentially associated with heterochromatin. Therefore, we concluded that *FRG2A* and its paralogs constitute a family of long non-coding RNAs associated with heterochromatin.

### Trans-interactions of FRG2A/B RNAs are enriched at highly repetitive loci: rDNA and centromeric satellites

Among *FRG2* paralogs, *FRG2A* is the only one that has been linked to a human disease, FSHD. The amount of FRG2A transcript (hereafter referred as FRG2A-t) is anomalously increased in cells and muscles from FSHD affected subjects (Figure 1b), and inversely correlates with the number of D4Z4 repeats at 4q35^17,20^. To investigate its function, we obtained a comprehensive FRG2A-t interactome by using ChIRP (Chromatin isolation by RNA purification) analysis^34,35^, which can simultaneously map RNA-RNA (ChIRP RNA-seq), RNA-protein (ChIRP-MS), and RNA-chromatin interactions (Figure 2a) by oligonucleotides-mediated purification of a target lncRNA. The experiment targeted both *FRG2A* (Chr4q35) and *FRG2B* (Chr10q26) transcripts (FRG2A/B-t) due to the 99.5% identity between the two genes (Figure 1a). This analysis uncovered a total of 2,313 genomic target sites for FRG2A/B-t, spanning 813,763 bp in HeLa cells (Figure 2b and Supplemental Table 3).

**Fig. 2.**
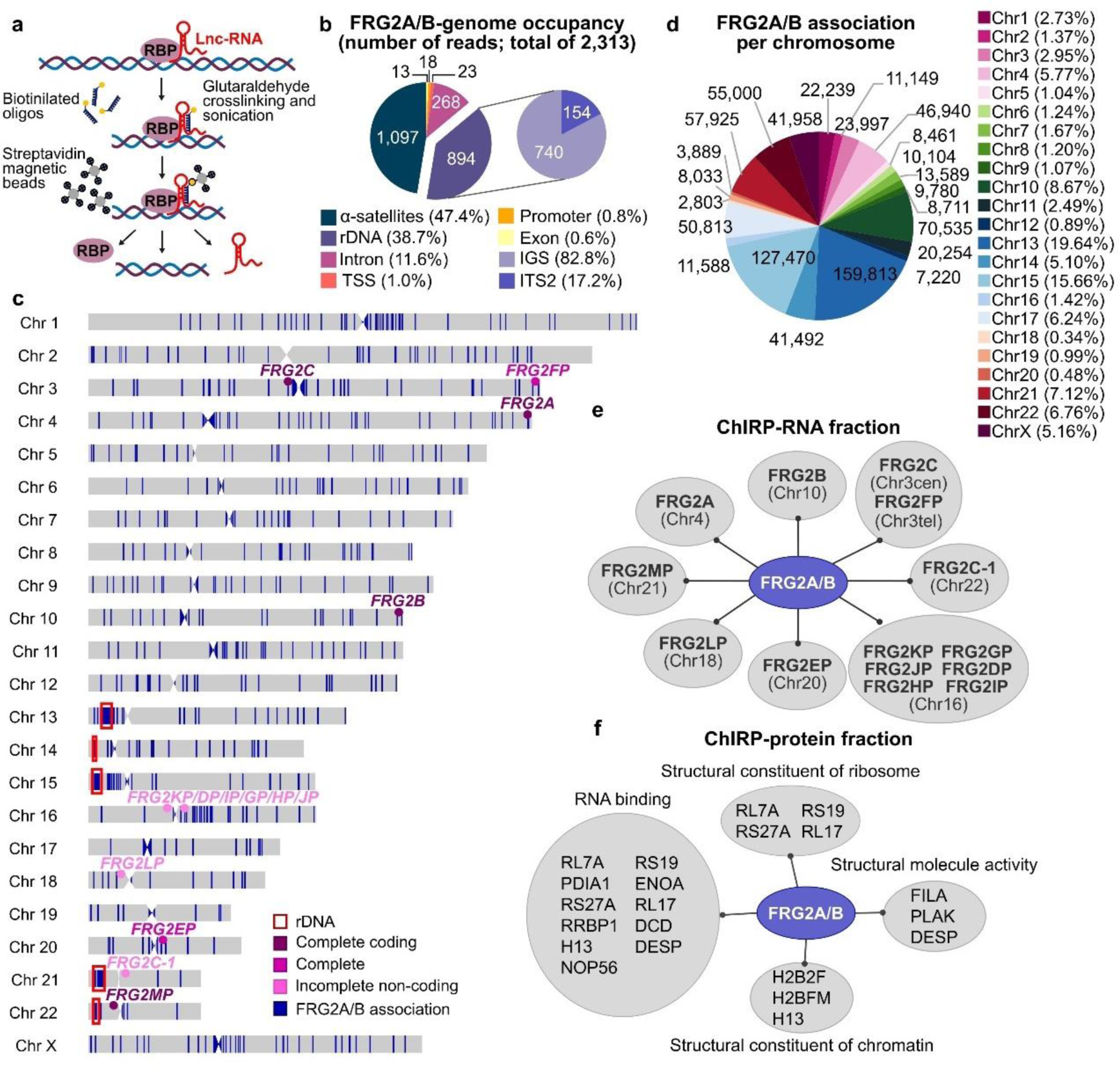
ChIRP assay revealed FRG2A/B interactome. **a**, Schematic diagram of the ChIRP workflow. **b**, Pie chart showing relative representation of various genomic regions obtained by the FRG2A/B ChIRP-DNA sequencing. A total of 2,313 genomic target sites were found, with the enrichment of highly repetitive elements clustered into two main regions: the centromeric α-satellites (47.4%) and the ribosomal DNA arrays (38.7%). Within rDNA arrays, FRG2A/B-t was found to be predominantly associated with the InterGenic Spacer (IGS) region (82.8%) and the ITS2 (17.2%). **c**, Chromosome ideograms overlapped with FRG2A/B-association track (in blue) displaying the enrichment of FRG2A/B-t at the centromeres and rDNA arrays. *FRG2* genes and rDNA arrays localization are indicated. **d**, Pie-chart showing FRG2A/B association, obtained from the ChIRP-DNA sequencing, per each chromosome. Values are in bp and percentages are indicated in the legend. **e**, FRG2A/B-interacting RNAs obtained by the ChIRP-RNA sequencing. **f**, FRG2A/B**-**interacting proteins subclassified into functional groups, acquired by mass-spectrometry analysis of the ChIRP-protein fraction

Interestingly, most of FRG2A-t-associated regions occurred with two types of repetitive elements: 47.4% of read coverage was from centromeric satellites and 38.7% from ribosomal DNA (rDNA) arrays (Figure 2b). Other types of repetitive regions such as telomeres or macrosatellites were not detected, nor were *in cis* interactions with the D4Z4 macrosatellite at the 4q35 and 10q26 subtelomeres. Hence. we concluded that FRG2A/B-t display specific localizations, preferentially associating *in trans* with the centromeres of all the human chromosomes, and within the rDNA arrays on all the acrocentric chromosomes as depicted in Figure 2c. We also observed a preferential (>5% of the total reads) association *in trans* with chromosome 17 and X centromeres and *in cis* with chromosome 4 and 10 centromeres (Figure 2d).

The specificity of FRG2A-t and FRG2B-t purification by ChIRP is demonstrated by the fact that of all the *FRG2* paralog genes, only the DNAs encoding these two transcripts were recovered in the bound fraction (Supplemental Table 3). In contrast, the ChIRP-RNA fraction obtained using the same oligonucleotide probes contained all FRG2s transcripts (FRG2s-t). This suggests that multiple distinct FRG2s-t could be associated and localized within the same trans-acting complexes, possibly in different combinations (Figure 2e and Supplemental Table 3).

### The FRG2A/B-t -interacting proteome

ChIRP-MS^36^ was also used to detect *FRG2*-associated proteins. ChIRP-protein samples were subjected to proteomic analysis through nanochromatography coupled to a high-resolution mass spectrometry. We averaged data from two replicate experiments which detected 36 enriched proteins (Supplemental Table 4). The list of proteins was analyzed by using the DAVID functional annotation tool and the GO_Molecular function annotation^37^, which revealed the enrichment of RNA-binding factors and structural components of ribosomes and chromatin (Figure 2f). Some of these proteins (RL7A, RS27A, RS19, RL17) are involved in ribosome biogenesis and homeostasis, which take place in the nucleolus.

### FRG2s RNA localize in the nucleolus

The ChIRP data raised the possibility that FRG2 RNAs could be linked to ribosome biogenesis and/or nucleolar function. We therefore assessed their localization via RNA-FISH experiments in primary myoblasts from control and FSHD samples. We used two different fluorescent LNA probes: one specifically recognizing FRG2A-t (LNA647-FRG2A) and one recognizing both FRG2A-t and FRG2B-t (LNATY563-FRG2A/B) (Figure 3a-f and Supplemental Figure 7a-j). Confocal imaging of LNATY563-FRG2A/B revealed the FRG2A/B-t signal was superimposable with the nucleolus both in control and FSHD cells (Figure 3a-f). To confirm the nucleolar localization of FRG2A/B-t, we performed RNA-FISH in Hela-KYOTO cells^38^, which express a nucleolin-GFP fusion protein to facilitate visualization of the nucleolus. Supplemental Figure 7k-n shows the co-staining between LNATY563-FRG2A/B and nucleolin-GFP. RNase A (Supplemental Figure 7o-r) and RNase H treatment (Supplemental Figure 7s-v) abolished the signal and confirmed that we were detecting FRG2A/B-t as DNA-RNA hybrids^39^. We concluded that FRG2A/B-t are enriched in the nucleolus.

**Fig. 3.**
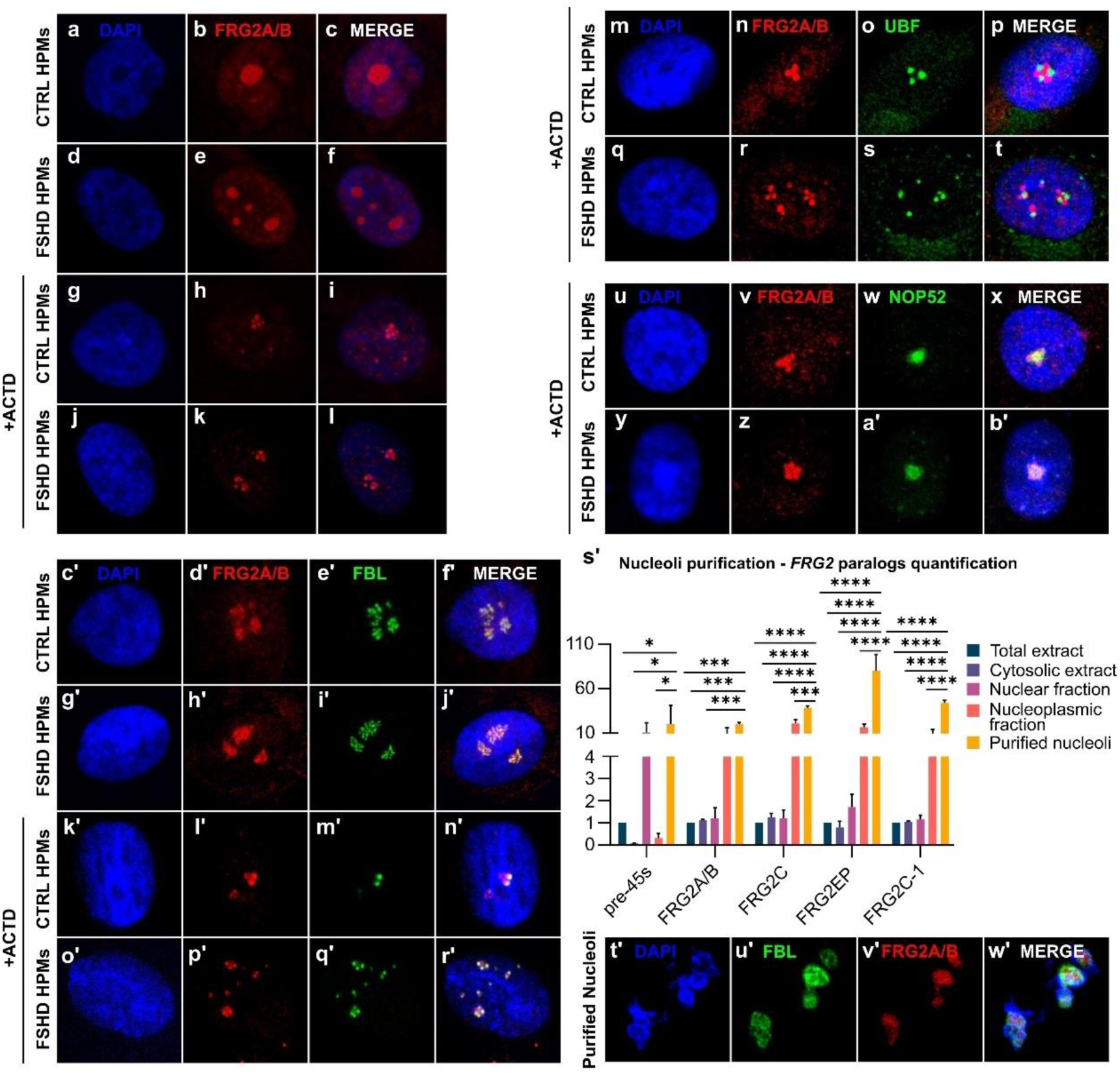
FRG2A/B is localized within the Dense Fibrillar Components of the nucleoli. **a-l**, RNA-FISH performed with FRG2A/B-specific fluorescent LNA probe (red), in human primary myoblasts (HPMs) obtained from CTRL and FSHD subjects. To induce nucleolar segregation, HPMs in panels g-l were treated with 1μg/mL of Actinomycin D (ACTD) for 4 hours. **m-b’,** Immuno-RNA-FISH performed with anti-UBF antibody (m-t) or anti-NOP52 antibody (u-b’) and FRG2A/B specific fluorescent LNA probe (red) in CTRL and FSHD-derived HPMs, upon ACTD treatment (1μg/mL for 4 hours). No colocalization was found between FRG2A/B dots and FC-UBF positive foci nor with GC-NOP52 positive domains. **c’-r’**, Immuno-RNA-FISH performed with anti-FBL antibody (green) and FRG2A/B specific fluorescent LNA probe (red) in CTRL and FSHD-derived HPMs, treated or not with ACTD (1μg/mL for 4 hours). FRG2A/B dots merged with DFC-FBL positive foci. **s’**, RT-qPCR of RNA fractions obtained from nucleoli purification experiment performed in HeLa cells. The analysis confirmed that FRG2A/B and its paralogs are enriched in the nucleolar fraction of the RNA. Statistical significance was tested by using two-way ANOVA test. p-values (****<0.0001) **t’-w’**, Immuno-RNA-FISH performed with anti-FBL antibody (green) and FRG2A/B-specific fluorescent LNA probe (red) in purified nucleoli isolated from HeLa cells, confirmed the colocalization between FBL and FRG2A/B.

To evaluate the precise localization of FRG2A/B-t within the nucleolus we performed immuno-RNA FISH in primary myoblasts using LNATY563-FRG2A/B and specific antibodies against UBF, FBL and NOP52 to visualize the Fibrillar Center (FC), Dense Fibrillar Component (DFC) and Granular Component (GC) regions respectively (Figure 3 and Supplemental Figure 7w-l’). We also performed these experiments after Actinomycin-D (ACTD)-induced nucleolar segregation^40–43^, which helps discriminating nucleolar compartments. In ACTD-treated cells, FRG2A/B-t staining was visible as discrete dots symmetrically organized around the nucleolus (Figure 3g-l and Supplemental Figure 7m’-x’) that were superimposable with nucleolar caps, usually appearing as one or more dots surrounding the central GC body.

Further, the co-staining experiments revealed that FRG2A/B-t did not co-localize with the UBF-positive FC compartment (Figure 3m-t), nor with NOP52-positive GC (Figure 3u-b’). In contrast, we detected the highest degree of overlap FBL and FRG2A/B-t, both in ACTD-treated and untreated cells (Figure 3c’-r’). These findings indicate that *FRG2*s-t are localized in the DFC nucleolar compartment, the site of rDNA transcription and very early ribosome biogenesis^44^. In these experiments, FSHD cells did not differ from control cells, showing comparable localization of nucleolar components and FRG2A/B-t.

To further validate our observations, we isolated nucleoli from HeLa cells^45,46^ (Supplemental Figure 8a). DNA and RNA were extracted from cytoplasmic, nuclear, nucleoplasmic and nucleolar fractions and the qPCR analysis of the DNA content in the different fractions was used to validate the purification of nucleoli (Supplemental Figure 8b, c). Analysis of the purified RNAs revealed that all FRG2s transcripts were enriched in the nucleolar-RNA fraction (Figure 3s’). Immuno-FISH experiments on isolated nucleoli from HeLa cells confirmed the presence of FRG2A/B-t and their colocalization with Fibrillarin (Figure 3t’-w’). These findings together with the ChIRP data and protein interactome data (Figure 2) indicate that FRG2s-t are nucleolar, localized in the DFC component and are sequestered in nucleolar caps during nucleolar segregation.

### FRG2A/B-t promote centromere-nucleolar associations in FSHD cells

The genomic association profile of FRG2A/B-t revealed frequent associations with rDNA arrays and with the centromeric regions of all chromosomes (Figure 2). In mammalian cells, these repetitive, heterochromatin regions are enriched around the periphery of nucleoli, as part of a subset of the genome known as Nucleolus Associated Domains (NADs)^47,48^.

To investigate the interaction between FRG2A/B-t and NADs, we first analyzed existing NADs mapping data^45^ from HeLa cells by using the NAD finder tool^46^ and the T2T-CHM13 genome assembly. Graphing the NADs (Supplemental Table 5, Figure 4 and Supplemental Figure 9) together with the genomic association profile of FRG2A/B-t obtained by ChIRP (Figure 2c), showed the extensive overlap between these two features (Figure 4a). Quantifying these data, we found that 86.3% of FRG2-associated regions overlap with NADs in HeLa cells (Figure 4b). The overlap included NADs on the rDNA-containing acrocentric chromosome arms (50.4%) and centromeric regions (49,6%) (Figure 4c, d).

**Fig. 4.**
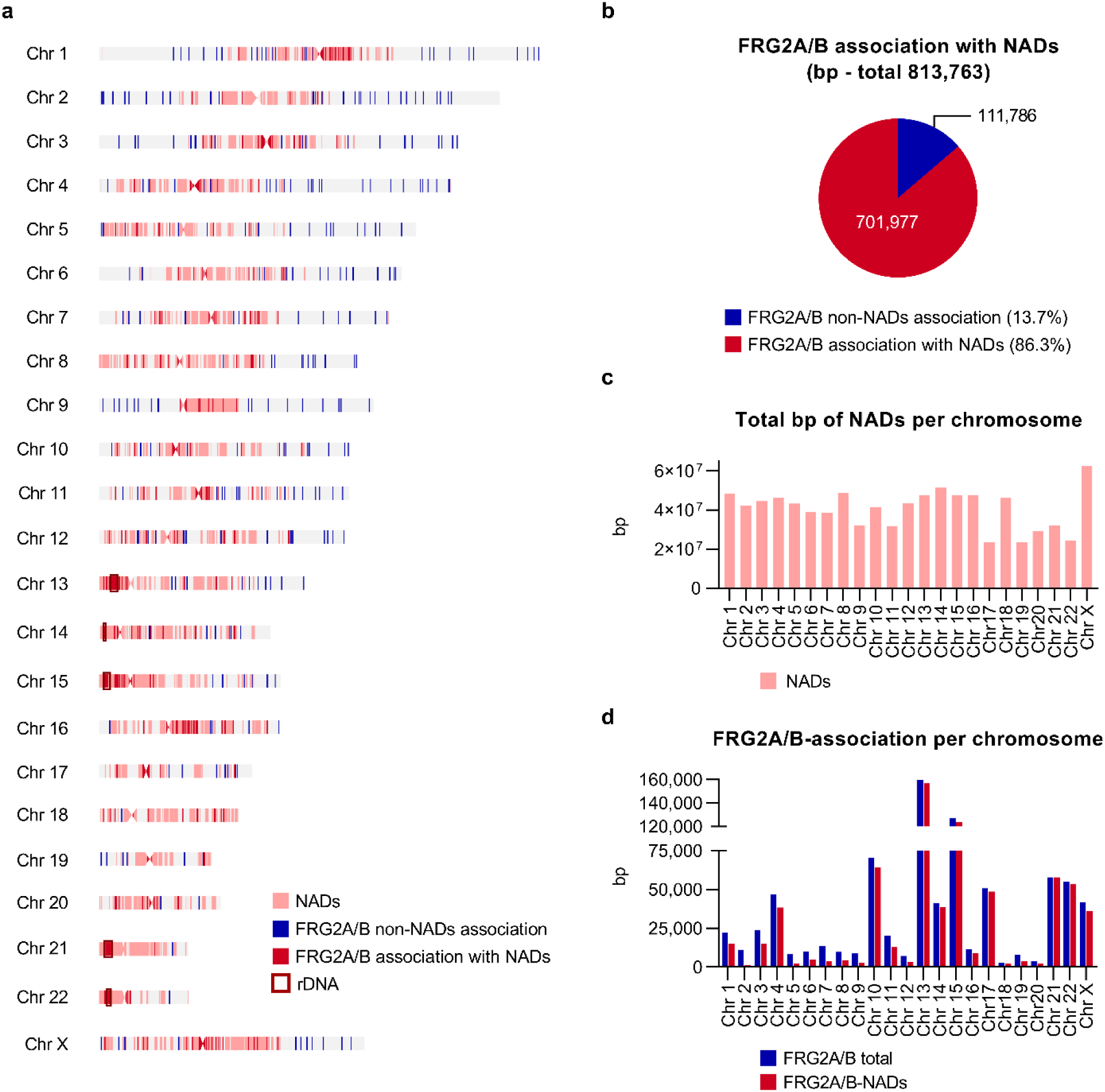
FRG2A/B-associated regions overlap NADs. **a**, Chromosome ideograms showing FRG2A/B-associated regions obtained by the CHIRP-DNA sequencing (blue) and NADs genomic regions (pink) extracted from existing data obtained from HeLa cells. The overlap between FRG2A/B and NADs tracks is shown in red. **b**, Pie chart showing the quantification of FRG2A/B-associated regions overlapping with NADs. **c**, Histogram displaying the distribution of NADs per chromosome (in bp), obtained from existing data in HeLa cells. **d**, Histogram showing FRG2A/B genomic association per chromosome (in bp). The total number of FRG2A/B-associated bp are shown in blue, and FRG2A/B-NADs bp are shown in red.

Within the centromeres, the ChIRP data indicated that the most frequent FRG2A/B-t associations occurred at the Higher Order Repeats (HORs) within α-satellites, which are the sites of kinetochore assembly and CENP-A/B enrichment at active centromeric repeat arrays^50^ (Figure 2 and Supplemental Figure 10).

These observations suggested that FRG2A/B-t may create a structural link between centromeres and nucleoli, thus regulating the nucleolus-associated chromatin interactions^48,49,51–52^. They also raised the question whether these chromosome features might be perturbed in FSHD patients.

To address these issues, we concomitantly investigated the centromeres and nucleoli localizations by immunofluorescence with antibodies raised against kinetochore protein CENP-B and the nucleolar FBL, in human primary cells (Figure 5a-h). We measured the distance between CENP-B foci and nucleoli (FBL staining for DFC), confirming a significant increase in the percentage of centromeres present at the nucleolar periphery in FSHD samples (Figure 5i). Instead, to confirm that this difference results from FRG2 overexpression rather than any other difference in genotype, we performed siRNA-mediated depletion of FRG2A/B-t in FSHD cells (Supplemental Figure 11) which reduced the frequency of centromere-nucleolus associations (Figure 5j-e’). This both confirmed a role for FRG2A/B in clustering centromeres at nucleoli and indicates that this is a dynamic process.

**Fig. 5.**
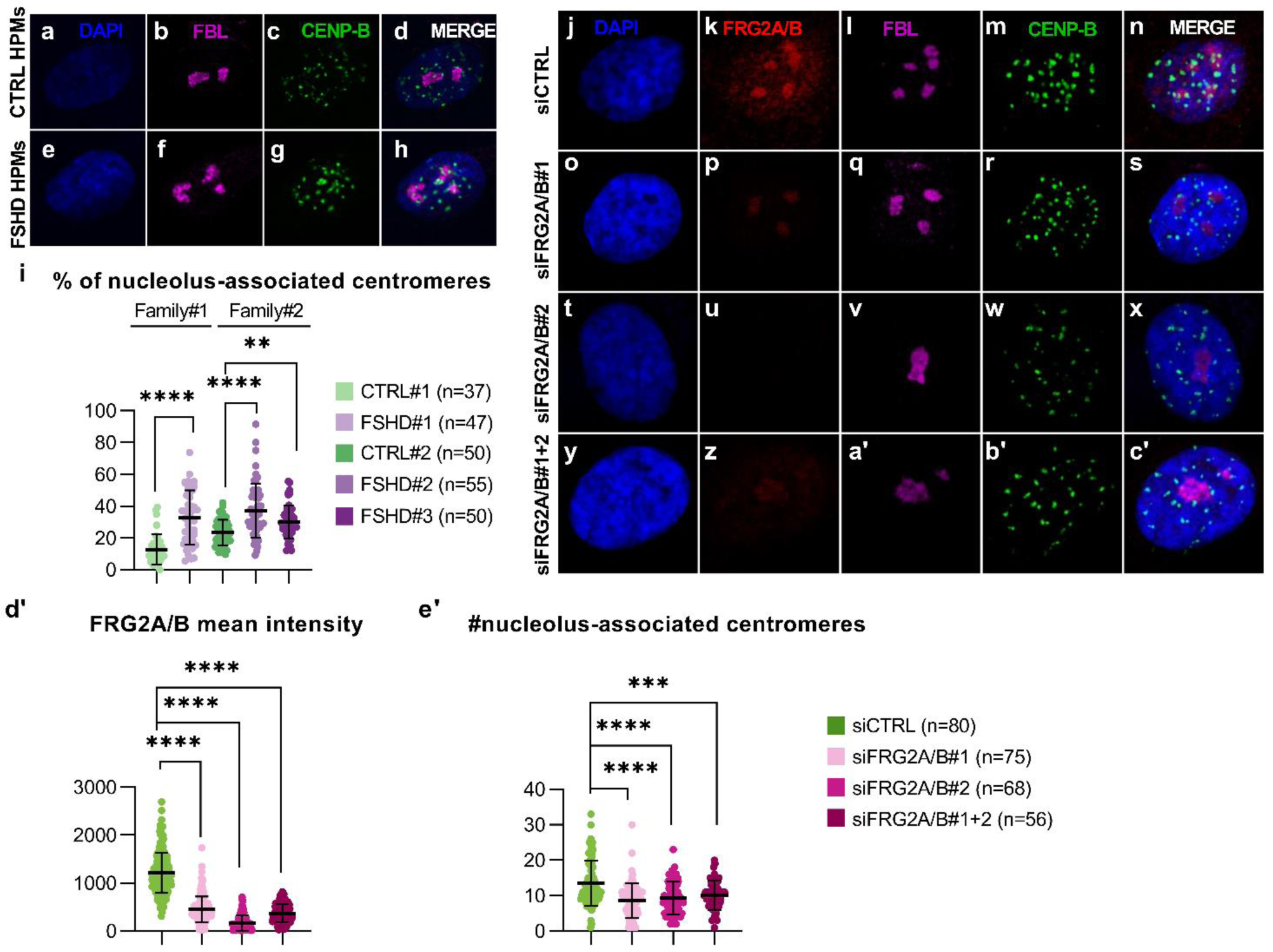
FRG2A mediates centromeres compaction at NADs in FSHD cells. **a-h**, Immunofluorescence assay performed with anti-FBL (magenta) and anti-CENP-B (green) antibodies in CTRL and FSHD-derived human primary myoblasts (HPMs). Increased association of centromeres with the nucleoli was detected in FSHD samples. **i**, Quantification of nucleolus-associated centromeres per cell from the experiment illustrated in panels a-h. Nucleolus-associated centromeres were defined as CENP-B-stained centromeres colocalizing with FBL, or in which FBL signal was within a range of 140 nm from CENP-B-positive foci. Analysis was conducted using ScanR software. Two families composed of CTRL and FSHD subjects were tested, and the number of cells analyzed per each subject is reported. Statistical significance was evaluated by using a Mann-Whitney test. p-values (****<0.0001). **j-c’**, Immuno-RNA-FISH experiments performed with anti-FBL (magenta) and anti-CENP-B (green) antibodies, and with FRG2A/B-specific fluorescent probe (red). The experiment was conducted in FSHD HPMs upon FRG2A/B silencing by two different siRNAs (siFRG2A/B#1 and siFRG2A/B#2), used alone and in combination (siFRG2A/B#1+2). A scrambled siRNA was used as a control (siCTRL). Upon FRG2A/B silencing, FRG2A/B fluorescent signal was reduced (**p**, **u**, **z**), and centromere localization was restored and visible as dotted staining scattered within the entire nuclei (**r**, **w**, **b’**). **d’**, Quantification of FRG2A/B mean intensity from panels j-c’. The analysis confirmed the reduction of FRG2A/B signal upon its siRNA-mediated silencing compared to siCTRL treatment. Quantification was conducted by using ScanR software, and statistical significance was tested by using two-way ANOVA test. p-values (****<0.0001). **e’**, Quantification of nucleolus-associated centromeres found in FRG2A/B-silenced FSHD HPMs, from panels j-c’. In siFRG2A/B treated cells nucleolus-associated centromeres are significantly reduced. Quantification was conducted by using ScanR software, and statistical significance was tested by using two-way ANOVA test. p-values (****<0.0001).

We concluded that FRG2A/B-t associates with rDNA repeats and centromeres, two main components of NADs. Furthermore, FRG2A/B-t promotes association of centromeres with the nucleolar periphery demonstrating an unforeseen mechanism for nuclear compartmentalization.

### The increased levels of FRG2A/B-t in FSHD cells impair nucleolar function

The localization of FRG2A/B-t within the DFC region of the nucleolus suggested a functional role(s) in rRNA transcription or processing^53^. To test this possibility, we first examined processing of rRNA transcripts in FSHD cells, but no defects were observed (Supplemental Figure 12), suggesting that post-transcriptional processing of rRNAs is not a regulatory target of FRG2 RNAs^53^. This is consistent with the observation that FRG2A/B is associated with the rDNA arrays (Figure 2c), but not with any rRNA species (Supplemental Table 3).

We therefore turned our attention to the potential roles of FRG2A/B-t in the regulation of rDNA transcription. Each unit of rDNA consists of the 13Kb transcribed region giving origin to the 45S precursor RNA (pre-45S), whose transcript is processed into mature 18S, 5.8S, and 28S RNAs, and a 30kb-long InterGenic Spacer (IGS), which is composed by small repetitive sequences^54^ (Supplemental Figure 13).

We analyzed the rDNA-associated ChIRP-reads to determine more precisely where FRG2A/B-t localize within human rDNA repeats. We found that 82.8% of rDNA ChIRP reads mapped to the IGS region, and 17.2% to the 5.8S 5’ region overlapping the Internal Transcribed Spacer 2 region (ITS2) (Figure 2b and Supplemental Figure 13). The FRG2A/B-t associated regions within the IGS span from IGS20/22 to IGS30/32, corresponding to CT-microsatellites (Supplemental Figure 13)^54–57^.

Several lncRNAs have been described to regulate rDNA transcription as well as to help in maintaining nucleolar structures and functions^58–63^. We thus reasoned that the association of FRG2A/B-t with IGS regions might have a similar role in modulating rDNA transcription as well. To explore this possibility, we measured steady-state levels of RNA Polymerase I-mediated transcription of rRNA by measuring the levels of pre-45S transcript in muscle samples and fibroblasts with similar genetic background, selecting samples from affected and unaffected FSHD siblings. As shown in Figure 6a and Supplemental Figure 14a, we observed significantly lower levels of pre-45S transcripts in FSHD myoblasts and not in fibroblasts which was paralleled by (Figure 6b) a significant enrichment in heterochromatin-associated histone mark H3me3K9 at IGS and at the rDNA promoter in FSHD cells in comparison with controls. This confirmed that muscle-specific FRG2s could be involved in in the regulation of rRNA transcription.

**Fig. 6.**
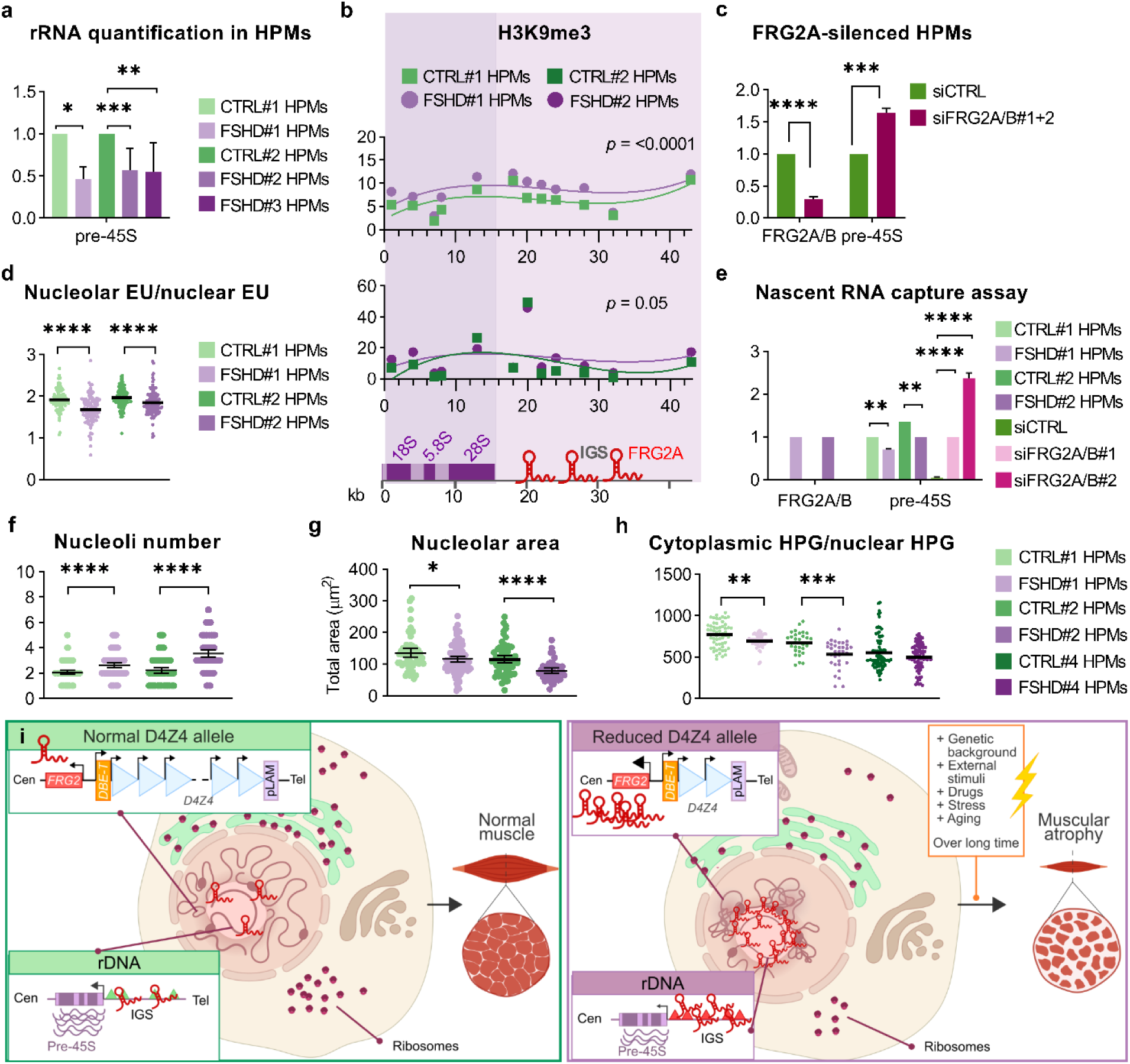
FRG2A/B increased levels impair nucleolar function in FSHD cells. **a**, RT–qPCR of HPMs cDNA of CTRL and FSHD subjects. Statistical significance was tested by the two-way ANOVA aalysis. p-value (****<0.0001). **b**, Chromatin-immunoprecipitation (ChIP) performed in CTRL and FSHD-derived HPMs showing H3K9me3 enrichment at IGS and rDNA promoter. Statistical significance (*p*=<0.0001 CTRL#1 vs FSHD#1 and *p*=0.05 CTRL#2 vs FSHD#2). A schematic representation of the rDNA array is reported below. **c**, RT-qPCR of FSHD HPMs cDNA upon FRG2A/B siRNA-mediated silencing. Statistical significance was tested by using the two-way ANOVA analysis. p-value (****<0.0001). **d**, Via 5-Ethynil-uridine (EU) incorporation, rRNA transcription was quantified as the ratio between nuclear/nucleolar EU fluorescent signals. Statistical significance was tested by using the Mann-Whitney test. **e**, RT-qPCR of nascent RNAs captured by EU incorporation in CTRL and FSHD HPMs, and upon siFRG2 treatments of FSHD-HPMs. Statistical analysis was performed by using the two-way ANOVA test. **f**-**g**, Nucleoli number and areas were measured in FSHD and CTRL HPMs. Statistical significance was tested by using the Mann-Whitney test. **h**, Protein synthesis levels in CTRL and FSHD HPMs by using <u>L-Homopropargylglycine</u> (HPG) incorporation. HPG signal was quantified as nuclear/cytoplasmic HPG ratio. Statistical significance was tested by using the Mann-Whitney test. p-values (***<0.001). **i**, A new multi-step mechanism for FSHD pathogenesis. In individuals carrying a normal-sized D4Z4 allele at 4q35, FRG2A is barely expressed and has no effects on genome architecture and rDNA arrays, allowing the normal growth and function of muscle. In subjects carrying a reduced D4Z4 allele (≤10 repats), FRG2A is aberrantly overexpressed. This causes the alteration of genome architecture and the impairment of rDNA transcription resulting in the reduction of protein synthesis, which, in the long term, may result in muscle atrophy.

To test the involvement of FRG2A/B-t in this process, we examined the effects of their siRNA-mediated depletion, observing increased levels of pre-45S transcripts both in HeLa cells and in human primary myoblasts derived from FSHD patient biopsies after silencing (Supplemental Figure 14b and Figure 6c).

To use a second approach to bolster the previous steady-state measurements, we pulse-labeled living cells with 5-ethynyl uridine (EU) followed by “click-it chemistry”^42,64,65^ to facilitate labeling and capture newly-synthesized RNAs. Nucleolar incorporation of the EU was quantified as the ratio of nuclear EU/nucleolar EU (Figure 6d and Supplemental Figure 15). Via biotinylation of EU labeled-newly-synthesized RNAs, we performed a nascent RNA capture assay quantified specific transcripts by RT-qPCR (Figure 6e). These experiments detected significantly lower levels of nascent pre-45S RNAs in FSHD samples compared to control samples.

Paralleling the steady state readout, the pre-45S levels were increased in FSHD cells upon silencing of FRG2A/B (Figure 6e). Furthermore, the assay confirmed that FRG2A/B are actively transcribed in myoblasts derived from FSHD patients but not healthy family members (Figure 6e). Altogether these results establish a previously unanticipated rRNA regulatory role for these lncRNAs. In addition to the transcriptional changes, we observed a significant increase in the average number of nucleoli per cell, and the reduction of singular nucleolar areas and of the total area of nucleoli per cell in FSHD cells (Figure 6 f,g). This is consistent with previous data indicating that the size of the nucleolus positively correlates with rRNA synthesis^66^ and that the downregulation of rRNA gene transcription leads to a reduction in nucleolar size^67^.

We hypothesized that these altered nucleolar characteristics would correlate with changes in protein synthesis capacity. To test this, we measured L-Homopropargylglycine (HPG)^68^ incorporation. Indeed, we found that the global rate of protein synthesis in FSHD cells (Figure 6h) was reduced compared with that of cells from their healthy relatives. Based on these findings, we conclude that the negative regulation of rRNA transcription imposed by excessive levels of FRG2A/B results in the impairment of protein synthesis in FSHD cells.

## DISCUSSION

Here, we discover a previously unrecognized family of regulatory lncRNAs. These FRG2 family members are localized to nucleoli and FRG2A is upregulated in FSHD patients, acting to reduce rRNA levels, thereby diminishing translational capacity.

Protein synthesis is a key molecular process for skeletal muscle maintenance and growth and requires ribosome biogenesis. However, ribosome biogenesis is reduced under catabolic states, in particular during decreased skeletal muscle contractile activity^69^. This suggests that maintenance above a certain threshold has the potential to reduce or prevent muscle atrophy during disuse, aging and disease^69^.

In the case of FSHD, the association of FRG2A/B-t with ribosomal DNA and its impact on protein synthesis, suggest a link between nucleolar dysfunction and the development of FSHD pathology. We have demonstrated that FRG2A-t is expressed in muscle cells and overexpressed in cells derived from FSHD patients. Therefore, conditions that increase levels of FRG2A/B-t are expected to impair ribosome biogenesis and protein synthesis, with negative effects on muscle functions eventually leading to disease^70^. Supporting this prediction, genotoxic agents and DNA damage are known to contribute to muscle wasting and aging when they persist over time^71,72^.

While the primary function of the nucleolus is ribosome biogenesis^73,74^, additional biological processes occur in nucleoli, for example the processing of the Signal Recognition Particle (SRP)^75^. It is also clear that the nucleolar periphery is an important hub for organizing the largely heterochromatin subset of the genome known as NADs^47^. Our studies show that FRG2 lncRNAs regulate interactions at the nucleolar periphery (NP) promoting centromere clustering and regulate rDNA transcription. Similarly, another lncRNA that contributes to chromosome anchoring to the NP is Firre, which is required for the inactivation of chromosome X (Xi)^76^. How lncRNAs regulate both nuclear architecture and gene expression are major questions for nuclear cell biology.

The discovery of the FRG2 lncRNA family is a foundational step towards a new paradigm in which repetitive elements exert trans-functions via lncRNAs. Noting that one contributing factor to FSHD disease is the deletion of D4Z4 macrosatellites near the FRG2A locus, we speculate that FRG2A represents just one example of repeat-regulated lncRNAs interacting in trans with many genomic loci and representing architectural and regulatory hubs in the mammalian cell nucleus.

Other lncRNAs play key roles in the formation of membrane-less nuclear bodies through Liquid-Liquid Phase Separation, serving as a scaffold to anchor proteins and mRNAs^77^, but in the previously studied cases there is a single locus encoding the lncRNAs. When encoded or regulated by repetitive loci, the potential for complexity escalates.

In this scenario, we speculate that the effects of FRG2A expression on translation might be affected not only by copy number variations of D4Z4 but also by the variable number of rDNA repeats. That is, the effects of FRG2 lncRNAs on rDNA or other TAR_ls_ targets could depend on the ratio of the numbers of each element. Indeed, we propose that the FRG2 lncRNAs not only contribute to FSHD disease phenotypes, but also provide a measure of the intra- and inter-familiar phenotypic variability associated with this disease. These ideas may propel the discovery of new roles for repetitive elements in cell biology, and also provide robust systems to test the effects of TAR_ls_ CNVs in shaping cell functions and their response to external stimuli.

## MATERIAL AND METHODS

### Cell culture

HeLa and HEK293 cells were cultured in Dulbecco’s Modified Eagle’s Medium (DMEM), supplemented with 10% fetal bovine serum (FBS), 1% glutamine and 1% penicillin-streptomycin. Trypsin-EDTA 1X in PBS was used to collect cells. Human chromosome hybrids (CHO/Hyb) were obtained from the Coriell Institute for Medical Research and maintained following the supplier’s instructions: chromosome 3/CHO hybrid (GM10253), chromosome 4/CHO hybrid (GM10115), chromosome 10 /CHO hybrid (GM10926) chromosome 20/CHO hybrid (GM13140) and Chromosome 22/CHO hybrid (GM10888). Control and FSHD-derived primary myoblasts were cultured in DMEM, supplemented with 20% FBS, 1% glutamine, 1% penicillin-streptomycin, 2 ng/mL epidermal growth factor (EGF) and 25 ng/mL of fibroblast growth factor (FGF). Trypsin 2.5% in HBSS, diluted 1:10 in PBS, was used to collect primary myoblasts. Myoblasts were purified form muscular biopsies (Supplemental Table1) as previously described^80^.

### FRG2 expression profiling with NGS

cDNA from human primary fibroblasts and human biopsies was used to amplify FRG2-exon4 with adapters-containing-pan-FRG2s primer pairs (Methods-Supplemental Table 1). PCR products were purified and subjected to NGSelect Amplicons service based on Illumina Platforms and provided by Eurofins Genomics. Read qualities were assessed by Fastqc^79^. Individual reports were merged by MultiQC ^80^and quality metrics were visually inspected.

Reads were aligned to the T2T human genome assembly using Bowtie2^81^. Sample-level alignment statistics were computed by samtools idxstats^82^. To evaluate the number of reads aligning to each paralog, we generated i. a BED file containing the genomic coordinates of the paralogs; ii. a BED file for each sample containing the genomic coordinates of read alignments, defined considering a window of 600 bp downstream the leftmost mapping position of the SAM files. For each sample we then compared the two sets of coordinates using bedtools intersect^83^ with -c option and reported the number of reads aligning to each paralog.

### CRISPR-Cas9 editing

Editing of HeLa and HEK293T cell lines was carried out using pSpCas9(BB)-2A-GFP (PX458) plasmid carrying Cas9 with 2A-EGFP and cloning backbone for sgRNA. To provide specificity to the editing, we followed one step digestion-ligation protocol supplied by the productor. gRNA spacer was designed using the CHOPCHOP^84^ web tool (Supplemental Table 6). Sense and antisense oligonucleotide carrying gRNA spacer targeting 5’ and 3’ end of *FRG2* were selected without paralog specificity. Single strand oligonucleotides with 40 bp homology arms were used as repair template to provide knock-in at *FRG2* region (listed in Supplemental Table6). Oligonucleotides were designed (IDT) with phosphodiester modification at 5’ and 3’ ends of the molecule to impair degradation. We transfected HeLa and HEK293T at 80% of confluency with Lipofectamine^TM^ 2000 (Invitrogen) following the supplier’s protocol. A total amount of 2 µg of plasmid DNA was transfected in each well of a 6 well tissue culture dish. Repair template was added to the transfection solution at the final concentration of 300 nM. Cells were incubated overnight with transfection medium, then medium was changed. 36 hours after transfection, cells were trypsinized and resuspended at 1×10^6^ cells/ml. Cells were sorted with FACS ARIA III cell sorter based on expression of EGFP tag on Cas9. Editing characterization of the 4 bulk populations was carried out with NGSelect Amplicons (Eurofins Genomics) on PCR products obtained by using adapters-containing primers set amplifying all FRG2’s paralogs (pan-edited FRG2s in Methods-Supplemental Table 1). Bioinformatic analysis was performed as previously described for FRG2 expression profiling with NGS. Design of primers and repair template used in each knock-in setting were displayed in Methods-Supplemental Table 1.

### RNA extraction and Real-time quantitative PCR (RT-qPCR)

Total cellular RNA was obtained from cell lines and HPMs by using PureLink RNA Mini Kit (Thermo Fisher Scientifc cat #12183018A), according to the manufacturer’s instructions. DNAse digestion and cDNA synthesis were performed by using Maxima H-cDNA Synthesis Master Mix, with dsDNase (Thermo Fisher M1682). Specific mRNA expression was assessed by qRTPCR (iTaq Universal SYBR® Green Supermix, BIORAD #1725120 in a CFX connect Real Time Machine BIORAD) using primers listed in Supplemental Table 6, normalized over RPLP0 and GAPDH housekeeping mRNAs.

### Protein extraction and Immunoblotting

Protein extraction was carried out starting from 5×10^5^ cells per sample. Cells were incubated in 50μL of extraction buffer (50 mM Tris-HCl, 400 mM NaCl, 1% NP-40, 1X Protease Inhibition Cocktail (supplier)) on an inverting wheel for 30 min in a cold chamber. After centrifugation 13000 rpm for 30’ at 4°C, supernatant fractions containing extracted protein were collected. Then, immunoblotting was performed to assess the presence of FRG2 proteins. Protein extracts were separated on 12% SDS-PAGE gel and then electrotransferred onto a nitrocellulose membrane, and subjected to overnight incubation with primary antibody, listed in Supplemental Table 6. Secondary antibodies conjugated with horseradish peroxidase were used and visualized by chemiluminescence using a ChemiDoc system (Bio-Rad).

### RNA fractionation

RNA fractionation was performed starting from 1×10^6^ cells. After collection, cells were lysed with 175 μl of cold cytoRNA solution (50 mM Tris HCl pH 8.0, 140 mM NaCl,1.5 mM MgCl_2_, 0,5% NP-40, 2mM Vanadyl Ribonucleoside Complex; Sigma) and incubated 5 min on ice. Cell suspensions were centrifuged at 4°C and 300xg for 2 min and the supernatant, corresponding to the cytoplasmic fraction, was transferred into a new tube and stored on ice. Pelleted nuclei were extracted with 175 μl of cold nucRNA solution (50 mM Tris HCl pH 8.0, 500 mM NaCl, 1.5 mM MgCl_2_, 0,5% NP-40, 2mM Vanadyl Ribonucleoside Complex; Sigma) and incubated 5 min on ice. The lysed nuclei were centrifuged at 4°C and 16,360xg for 2 min and the supernatant, corresponding to the nuclear-soluble fraction, was transferred into a new tube and stored in ice. The remaining pellet was collected corresponding to the chromatin-associated fraction. Total RNA from the cytoplasmic and nuclear fractions was extracted by using PureLink RNA MiniKit (Invitrogen) following the manufacturer’s instructions for the RNA extraction from aqueous solutions. The pellet containing the chromatin-associated fraction was extracted with the standard procedure. cDNAs were obtained as described above.

### Chromatin-RNA immunoprecipitation (ChRIP)

Chromatin RNA immunoprecipitation (ChRIP) was performed as described in the published protocol^33^ using antibodies anti-H3K4me3, H3K9me3 H3K27me3, as reported in Methods-Supplemental Table 1. 3 × 10^6^ HPMs were used for each IP. RNA was extracted and RT-qPCR performed as described above. Ten percent input was used to calculate the percentage of transcript bound to chromatin compared to the negative control IgG.

### Chromatin Isolation by RNA purification (ChIRP)

ChIRP was performed in HeLa cells by using the Magna ChIRP^TM^ RNA Interactome Kit (Sigma, cat #17-10494) strictly following manufacturer’s instructions. We designed targeted biotinylated antisense oligonucleotide probes as shown in Supplemental Figure 16, complementary to the predicted FRG2A mature transcript (NMNM_001286820.2) by using Stellaris (ChIRP) Probe Designer tool, generating a custom ChIRP probe set consisting of 10 specific antisense oligonucleotides. ChIRP probes were divided in two sets: (i) FRG2 EVEN, containing 5 out of 10 antisense oligonucleotides (sets 2, 4, 6, 8, 10), and (ii) FRG2 ODD, containing the remaining 5 antisense oligonucleotides (sets 1, 3, 5, 7, 9). To rule out artefacts due to direct probes hybridization to genomic DNA, we also used two controls: (iii) LacZ probe set, which would pull down potential contaminating DNA and (iv) TERC probe set, which was also used as a positive control for the effective success of the immunoprecipitation protocol. The isolated RNA and DNA samples were then subjected to Next Generation Sequencing, while the protein fraction was subjected to mass-spectrometry analysis.

### ChIRP Bioinformatic Analysis

ODD and EVEN sequencing data were initially analysed in parallel by applying the same workflow. Raw data were trimmed by using fastp^85^ (-q 20 -u 20 -l 25 -g 10 -w 15) to remove low quality sequences. Retained high quality reads were mapped against the T2T human genome^86^ by using bowtie2 (--no-unal --no-discordant). By using samtools^87^, the obtained SAM files were binary compressed in BAM (view -bS) and sorted. Optical and PCR duplicates were marked by using PICARD^88^ MarkDuplicates (--REMOVE_DUPLICATES false --ASSUME_SORT_ORDER coordinate). Following ODD and EVEN alignment data per each experiment, were intersected by using the bedtools^89^ function intersect (-wa) in both direction (i.e. ODD to EVEN and EVEN to ODD). The obtained bam files were sorted and merged by samtools. Finally, merged bam files were sorted and indexed by samtools. Peaks calling was performed by using the macs3^90^ callpeak function(-f BAMPE -g 3.05e+9 -B -q 0.01). Regardless the analysed experiment, peak calling was performed by using input genomic DNA as control. Finally, the annotatePeaks.pl Perl script, belonging to the Homer suite^91^, was used to deduce annotation associated to the genomic regions in which peaks fell. Raw data and processed files for the ChIRP sequencing have been deposited in the NCBI Gene Expression Omnibus under accession number: PRJNA1122505.

### NADs annotation NAD finder Overlap

Sequencing data belonging to the Bioproject PRJNA354332^46^ and regarding WGS and Nucleolus Sequencing (NS) in untreated HeLa cell and relative replicates were retrieved from the SRA repository by using the sratoolkit prefetch and fastq-dump tools (--split-files --gzip). Raw data were trimmed by using fastp^85^ (-q 20 -u 20 -l 25 -g 10 -w 15) to remove low quality sequences. Retained high quality reads were mapped against the T2T human genome^86^ by using bowtie2 (--no-unal). By using samtools^87^, the obtained sam files were binary compressed (view -bS), sorted (sort) and, indexed (index). NS replicates were used by the R library HMMtBroadPeak (binsize=50,000, background=25, pseudocount=1) to identify NADs (Nucleolus Associated Domains) using WGS data as control. HMMtBroadPeak, relies on the same strategy implemented in NADfinder^92^, but it is optimized to overcome memory leaps. In particular, per each bin the number of reads falling within is counted and bins supported by a numbered reads at least equal to then background parameter are retained and normalized to CPM (count per million). Then log2 transform is applied to the ratio among NS and WGS normalized counts. Transformed values are finally used to define NAD peaks by running the Hidden Markov Model implemented in the R HMMt library^93^.

Overlaps between FRG2 peaks and inferred NADs were identified by using the bedtools intersect function (-wa -f). In particular, an increasing value of the required overlap (1 bp, 50% and 90% peak region) among peaks and NAD for reporting, was applied to reduce the effect of casual co-localization. Moreover in order to further evaluate the non-causality of peaks-NAD co-localization, 10,000 simulations of randomly extracted regions were produced by using bedtools random and the overlaps were measured as previously described. Finally, the R function ppois was applied to infer the probability to observe a number of overlaps greater than the one observed between FRG2 peaks and NADs, using the Poisson cumulative density function.

### ChIRP-MS

ChIRP isolated proteins were subjected to reduction with DTT 200 mM, to alkylation with IAM 200mM and to complete protein digestion with 1 μg of Trypsin (Sigma-Aldrich Inc., St. Louis, MO, USA). The peptide digests were desalted on the Discovery® DSC-18 solid phase extraction (SPE) 96-well plate (25 mg/well) (Sigma-Aldrich Inc., St. Louis, MO, USA), and after the desalting process, the sample was vacuum-evaporated and reconstituted in mobile phase for the analysis. The digested peptides were analyzed with an EASY nano-LC 1200 system (Thermo Scientific, Milano, Italy) coupled to a 5600+ TripleTOF system (AB Sciex, Concord, Canada). The liquid chromatography parameters were as follows: analytical column Acclaim PepMap C18 2μm 75µm x 150mm and injection volume 2 μL. The flow rate was 300 nL/min, phase A was 0.1% formic acid/water and phase B was 80% acetonitrile/0.1% formic acid/20% water. A two hours gradient was used (3-45%). For identification purposes the mass spectrometer analysis was performed using a mass range of 100–1600 Da (TOF scan with an accumulation time of 0.25 s), followed by a MS/MS product ion scan from 400 to 1250 Da (accumulation time of 5.0 ms) with the abundance threshold set at 30 cps (40 candidate ions can be monitored during every cycle). The ion source parameters in electrospray positive mode were set as follows: curtain gas (N2) at 30 psig, nebulizer gas GAS1 at 25 psig, ionspray floating voltage (ISFV) at 2700 V, source temperature at 90 °C and declustering potential at 85V. The mass spectrometry files were searched using Mascot v. 2.4 (Matrix Science Inc., Boston, USA) with trypsin as digestion enzyme, with 2 missed cleavages and a search tolerance of 50 ppm was specified for the peptide mass tolerance, and 0.1 Da for the MS/MS tolerance. The charges of the peptides to search for were set to 2 +, 3 + and 4 +, and the search was set on monoisotopic mass. The instrument was set to ESI-QUAD-TOF and the following modifications were specified for the search: carbamidomethyl cysteines as fixed modification and oxidized methionine as variable modification. The UniProt Swiss-Prot reviewed database containing human proteins (version 01/02/2018, containing 42271 sequence entries) was used.

### HeLa cell proteomics

After cell lysis with RIPA buffer, proteins were digested with trypsin and peptides were analyzed on an Ultimate 3000 RSLC nano coupled directly to an Orbitrap Exploris 480 with a High-Field Asymmetric Waveform Ion Mobility Spectrometry System (FAIMS) (all Thermo Fisher Scientific). Samples were injected onto a reversed-phase C18 column (15 cm × 75 µm i.d., Thermo Fisher Scientific) and eluted with a gradient of 6% to 95% mobile phase B over 41 min by applying a flow rate of 500 nL/min, followed by an equilibration with 6% mobile phase B for 1 min. The acquisition time of one sample was 41 min and the total recording of the MS spectra was carried out in positive resolution with a high voltage of 2500 V and the FAIMS interface in standard resolution, with a CV of −45V. The acquisition was performed in data-independent mode (DIA): precursor mass range was set between 400 and 900, isolation window of 8 m/z, window overlap of 1 m/z, HCD collision energy of 27%, orbitrap resolution of 30000 and RF Lens at 50%. The normalized AGC target was set to 1000, the maximum injection time was 25 ms, and microscan was 1. For DIA data processing, DIA-NN (version 1.8.1) was used: the identification was performed with “library-free search” and “deep learning-based spectra, RTs and IMs prediction” enabled. Enzyme was set to Trypsin/P, precursors of charge state 1–4, peptide lengths 7–30 and precursor m/z 400– 900 were considered with maximum two missed cleavages. Carbamidomethylation on C was set as fixed modification and Oxidation on M was set as variable modification, using a maximum of two variable modifications per peptide. FDR was set to 1%.

### Riboseq data analysis

First, we accessed (April 2024) the RiboSeq data portal^94^ to select RiboSeq experiments exploiting HeLa cells (Toggles: Available in Trips-Viz,Organism: Homo sapiens, Library-Type: Ribo-Seq, Inibitor: None). Following considering the selected samples, RiboSeq profiles for ENST00000226798 (FRG1 gene), ENST00000504750 (FRG2 gene), ENST00000443774 (FRG2B gene) and, ENST00000308062 (FRG2C gene) transcripts were extracted as CSV files from Trips-Viz^95,96^. Moreover, the RiboSeq profiles of ENST00000395240 (ALDOA gene), ENST00000501122 (NEAT1 gene) transcripts were also accessed as example of protein coding and non-coding genes, respectively.

### RNA-FISH and immunofluorescent-RNA-FISH (IF-RNA-FISH)

RNA-FISH experiments were performed as previously described^97^ in HeLa cells and human primary myoblasts by using Locked Nucleic Acid (LNA) fluorescent probes (FRG2A/B-LNATY563 and FRG2A-LNA665), designed by Genglobe LNA designer (Qiagen). Probes sequences and localization are shown in Methods-Supplemental Table 1. Where indicated, cells were treated with 1 μg/mL Actinomycin D (ActD) for 1-4 hours. Briefly, cells were fixed with 3% paraformaldehyde (PFA) in PBS, for 12 min at room temperature, and permeabilized with 0.5% Triton X-100 in PBS. As controls, RNase A or H treatments were also performed by incubating the slides with 1mg/mL RNase A/H in PBS at room temperature for 30 min. Slides were then stored in 70% ethanol (EtOH), at −20°C for at least one night. Next day, slides were dehydrated in 80%, 95% and 100% EtOH, 3 min each and left to dry. Hybridization solution was prepared as follow: 0.2 μM LNA probe, 50% formamide, 2X saline sodium citrate (SSC), 10% dextran sulphate, 10 mM vanadyl ribonucleoside complex (VRC), 2 mg/mL bovine serum albumin (BSA). The mix was denatured at 75°C for 7 min. Hybridization was performed in a humid chamber for 35 min at 37°C, protected from light. Coverslips were washed three times in 0.1X SSC heated at 60°C for 5 minutes, rinsed in 2X SSC and mounted by using the ProLong^TM^ Diamond Antifade Mountant with DAPI (Invitrogen™, cat #P36962).

For immunofluorescence-RNA-FISH (IF-RNA-FISH), slides were first subjected to IF protocol as follows. Coverslips were incubated in blocking solution (1% BSA in PBS) for 15 min, and then with primary antibodies (diluted 1:100) in blocking solution for 2 hours at room temperature. After washes in PBS, the cells were incubated with the secondary antibodies, diluted 1:250 in blocking solution, for 45 min. Slides were post-fixed in 3% PFA/PBS for 10 min at room temperature, washed in 2X SSC and then RNA-FISH protocol was followed as described above. A list of the antibodies used in this paper is provided in Methods-Supplemental Table 1.

Images of RNA-FISH and IF-RNA-FISH experiments were acquired by using the Leica TCS SP8 AOBS confocal microscope system equipped with a 63X oil immersion lens. Emission wavelengths were: 405 nm for DAPI signal, 488 nm for green staining, 550 nm for red fluorescence and 647 nm for magenta staining.

### Nucleoli isolation

Nucleoli were isolated from HeLa cells as previously described^46^ with some variations. Briefly, 1×10^8^ HeLa cells were washed three times with cold PBS. Cells were scraped and collected by centrifugation at 218xg and 4°C for 5 min. Cells were subsequently resuspended in 5 ml of cold Buffer A (10 mM Hepes pH 7.9, 10 mM KCl, 1.5 mMMgCl2, 0.5 mM DTT, EDTA-free protease inhibitors, Roche) and incubated 15 min on ice. Cells were then dounce homogenised 20 times using a tight pestle and centrifuged at 218xg and 4°C for 5 min. The supernatant was collected as cytoplasmic fraction, while the pellet, which contained nuclei, was resuspended with 3 ml of S1 solution (0.25 M sucrose, 10 mM MgCl_2_, EDTA-free protease inhibitors). Resuspended nuclei were layered over 3 ml of S2 solution(0.35 M sucrose, 0.5 mM M MgCl_2_, EDTA-free protease inhibitors) and centrifuged at 1,430g and 4°C for 5 min. The resulting nuclear pellet was resuspended with 3 ml of S2 solution and 10-sec bursts (with 10-sec intervals between each burst) using Soniprep 150 (MSE) with a fine probe sonicator and set at power setting 5. The sonicated sample was layered over 3 ml of S3 solution (0.88 M sucrose, 0.5 mM MgCl_2_, EDTA-free protease inhibitors) and centrifuged at 3,000g and 4°C for 10 min. The supernatant was collected as nucleoplasmic fraction. The pellet, which contained nucleoli, was resuspended in 0.5 ml of S2 solution, centrifuged at 1,430g and 4°Cfor 5 min, resuspended in 0.1 ml of S2 solution, and used to assess nucleoli purity. DNA and RNA were extracted from each fraction by using DNeasy® Blood and Tissue Kit (QIAGEN, cat#69504) and Pure Link RNA Mini Kit (Invitrogen™ cat #12183018A), respectively. Enrichment of target DNA sequences and RNAs were assessed by RT-qPCR, as previously described, using primers listed in Methods-Supplemental Table 1.

### Sequence of Primers and Probes

The sequences of the primers and LNA fluorescent probes used in this study are provided in Methods-Supplemental Table 1.

### Short-interfering RNA gene silencing

FRG2A/B transcripts were silenced taking advantage of short interfering RNAs (siRNA). We designed two FRG2A/B-specific siRNAs (Supplemental Figure 16 for sequences and localization) and silencer select siRNA (Invitrogen Silencer™ Select Pre-Designed siRNA Catalog number:4392420) and a scramble siRNA was used as a control. Lipofectamine™ RNAiMAX Transfection Reagent (Invitrogen, cat #13778075) was diluted in Opti-MEM™ I Reduced Serum Medium (Gibco™, cat #31985062). CTRL and FRG2A/B-specific siRNAs were diluted in Opti-MEM™ I Reduced Serum Medium, at 20 pM final concentration. Diluted siRNAs and Lipofectamine™ RNAiMAX Transfection Reagent were mixed together and incubated at room temperature for 15 min, to allow complexes to form. StealthTM FRG2A/B/control siRNA - Lipofectamine™ RNAiMAX Transfection Reagent complexes were added to each well containing cells and medium and incubated at 37°C in a humidified CO_2_ incubator for 48 hours.

### Northern Blot

Ribosomal RNA maturation pathways were evaluated by northern blotting, as described^98^. Briefly, 4 µg of total RNA extracted from HeLa cells and HPMs was diluted in 5 volumes of Glyoxal mix (6mL DMSO, 2 mL deionized glyoxal (Fluka, cat. #50649), 1.2 mL 10X BPTE, 600 µL 80% glycerol, 40 µL 10 mg/mL EtBr), incubated at 55°C for 1 hour and chilled on ice for 10 min. Then, an electrophoresis run was performed in a 1.2% agarose gel prepared in BPTE buffer (for 10X concentration: 30 g of 100 mM PIPES (SIGMA, cat. #P6757), 60 g of 300 mM BIS-TRIS (SIGMA, cat. #B9754), 20 mL 0.5 M EDTA pH 8.0), for 5 hours at 100 V. When migration was sufficient, the gel was rinsed twice with H_2_O, and soaked for 20 min in 75 mM NaOH and twice in 0.5 M Tris-HCl pH 7.4, 1.5 M NaCl for 10 min. RNA was transferred to a Hybond N+ (GE) membrane, O/N at room temperature. After a pre-hybridization step with tRNA from E. coli, hybridization was performed by using ^32^P-labelled ITS1 and ITS2 specific probes, incubated O/N at 42°C. Membranes were washed three times in 2X SCC + 0.1% SDS an twice in 0.5X SCC + 0.1% SDS. Radioactive signals were detected by using Typhoon Trio PhosphorImager and quantified using the MultiGauge software.

### Chromatin immunoprecipitation (ChIP)

Chromatin immunoprecipitation (ChIP) assays were performed in HPMs as described earlier in^99^ using specific antibodies, as listed in Methods-Supplemental Table 1. Enrichments of DNA sequences inimmunoprecipitates from at least three independent ChIP results were analyzed by quantitative real-time PCR (qPCR) as described above. Enrichment of amplified DNA sequences (primers listed in Methods-Supplemental Table 1) in immunoprecipitates was calculated as the ratio between the DNA amount in immunoprecipitation samples and that in the total input chromatin.

### Nascent RNA detection

Global RNA synthesis in CTRL and FSHD HPMs was visualized and quantified by using Click-iT™ RNA Alexa Fluor™ 488 Imaging Kit (Invitrogen™, cat #C10329). This technology exploits the incorporation of the uridine analogue 5-Ethynyl uridine (EU) within nascent RNA transcripts. Then, through a click-it chemistry, Alexa Fluor™ 488 fluorophore was bound to the incorporated EU, to temporal and spatial visualize the newly synthesized transcripts.

Briefly, the day before, HPMs were plated into a multi-24 wells plate (2×10^5^ cells per well), with 12 mm ⌀ coverslips placed at the bottom. Cells were incubated 1 hour with EU at 1 mM final concentration, under normal cell culture conditions. After, cells were fixed in 3.7% formaldehyde in PBS and permeabilized with 0.5% Triton X-100 in PBS. The Click-iT® reaction was carried out following the manufacturer’s instructions. Then, since the ScanR software required labeled nucleoli to quantify EU nucleolar signals, immunofluorescence with anti-fibrillarin antibody was performed, as previously described.

### Nascent RNA capture assay

Newly synthesized RNA in HPMs was also measured by using the Click-iT™ Nascent RNA Capture Kit (Invitrogen™, cat #C10365), another technology based on EU incorporation. Total RNA or mRNA labeled with EU is isolated and used in a click-it reaction, which binds a biotin azide to the incorporated EU, creating a biotin-based handle for capturing nascent RNA transcripts on streptavidin magnetic beads.

The day before, HPMs were plated into a T75 flask. The day after, living cells were incubated with 0.2 mM EU for 1 hour, under normal cell culture conditions. Then, total RNA was extracted as described above. EU-labeled RNA was biotinylated through the click-it reaction with 0.5 mM of biotin azide, following the manufacturer’s instructions. EU-biotin-labeled RNA was isolated by using Dynabeads® MyOne™ Streptavidin T1 magnetic beads and retro-transcribed as previously described. The cDNAs obtained were quantified by RT-qPCR, to assess the amount of rRNAs with primers listed in Methods-Supplemental Table 1.

This experiment was performed both in CTRL and FSHD HPMs, as well as in FRG2A/B-silenced FSHD HPMs, treated with siRNAs 48 h before EU incorporation, as previously reported.

### Nascent Protein synthesis detection

Nascent protein synthesis was analyzed in HPMs by using the Click-iT™ HPG Alexa Fluor™ 488 Protein Synthesis Assay Kit (Invitrogen™, cat #C10428). Click-iT® HPG (L-homopropargylglycine) is an amino acid analog of methionine, which is incorporated into proteins during active protein synthesis. Detection of the incorporated amino acid is allowed by a click-it reaction with Alexa Fluor® 488.

The day before, HPMs were plated into a multi-24 wells plate (2×10^5^ cells per well), with 12 mm ⌀ coverslips placed at the bottom. Cells were incubated with HPG at 50 μM final concentration for 30 min, under normal cell culture conditions. After, cells were fixed in 3.7% formaldehyde in PBS and permeabilized with 0.5% Triton X-100 in PBS. The Click-iT® reaction was carried on following the manufacturer’s instructions.

### High content imaging-based assa**y**

For these analyses, images were obtained using a Nikon Confocal Microscope A1 equipped with a 60X oil immersion lens. FRG2A/B foci, FBL staining, CENP-B foci, EU staining, nuclear/nucleolar EU ratio, HPG staining and nuclear/cytoplasmic HPG ratio were analysed using the ScanR Analysis software (Olympus).

To count the nucleolus-associated centromeres in HPMs, each centromere was segmented based on CENP-B signal, using an edge detection algorithm. Following segmentation, all CENP-B foci were counted and those colocalizing with FBL or in which FBL was within a range of 140 nm were considered as nucleolus-associated centromeres. The analysis was performed in CTRL and FSHD HPMs, as well as in FSHD HPMs after FRG2A/B siRNA treatment. In the latter, we also measure FRG2A/B mean intensity within the nucleoli as the ratio between the mean fluorescence intensity inside and around the nucleolus.

Images from the Click-iT™ RNA Alexa Fluor™ 488 Imaging Kit assay and Click-iT™ HPG Alexa Fluor™ 488 Protein Synthesis Assay Kit followed a similar analysis. To quantify EU incorporation within the nucleoli as a measure of rRNA transcription, nucleoli were segmented based on FBL signal. EU incorporation was quantified as the ratio between the EU signal inside the nucleolus and from the surrounding nuclear region. To quantify HPG incorporation, nuclei were segmented based on DAPI fluorescent signal. Then, HPG was measured as the ratio between the HPG signal from the cytoplasm and the nucleus.

All these data were plotted as box plots. Statistical significance was tested by using the Mann-Whitney test, when comparing CTRLvsFSHD HPMs, and two-way ANOVA for siRNAs treatments.

## Author Contributions

VS and FL were involved in the study design, molecular analysis, data collection, data analysis and interpretation, literature review, and the preparation of figures, tables, and manuscript writing. BF and SP conducted the bioinformatic analyses. MF contributed to the design and analysis of the CRISPR-Cas9 studies. MM carried out the proteomic analyses. GV, TM, and LM were responsible for patient clinical assessments and sample collection from FSHD families. AKH investigated ribosome biogenesis in patients. GP and PDK contributed to data interpretation and manuscript editing. BMS participated in the study design, data interpretation, and manuscript writing. RT coordinated the study design, data interpretation, and manuscript writing.

## Acknowledgements

We are indebted to all patients and their families for their participation in this study. We thank Jhonathan Vinet for technical assistance and imaging; Serena Carra for providing reagents; Donata Orioli and Kathleen Collins for their comments and assistance in editing the manuscript, and Matteo Chiara for his bioinformatic training to SP. We extend our gratitude to Professor Michael R. Green for his contributions to our research, and we honor his memory.

## Data Availability

Correspondence and requests for materials should be addressed to Prof. Rossella Tupler.The datasets generated during and/or analyzed during the current study are available in the NIH repository, BioProject: PRJNA1122505 and ProteomeXchange Consortium via the PRIDE partner repository with the dataset identifier PXD053349.

## SUPPLEMENTAL FIGURES

**Supplemental Figure 1.**
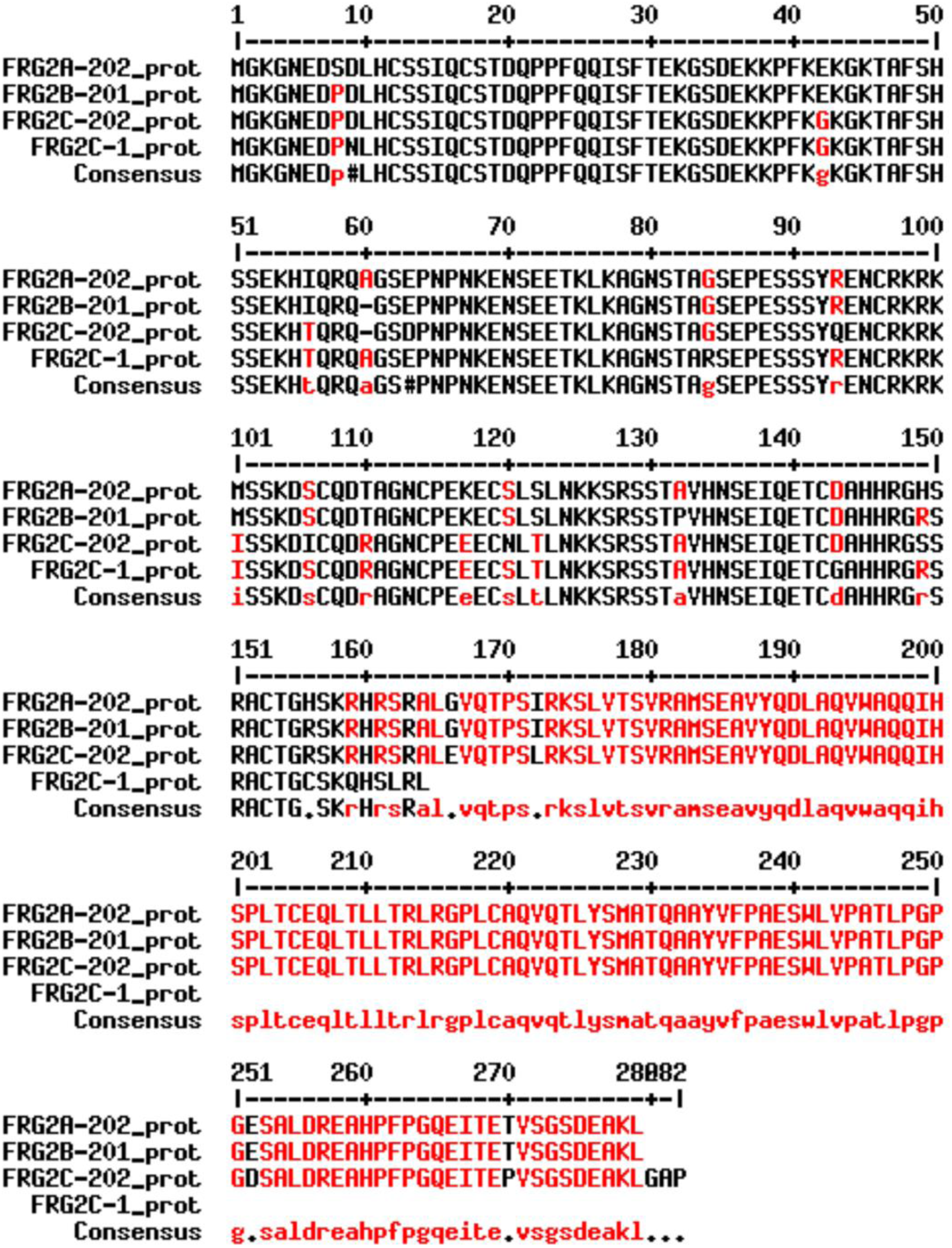
Sequences of the predicted FRG2 proteins. Alignment of the amino acid sequences of the predicted four FRG2 protein paralogs (FRG2A from chr 4, FRG2B from chr 10, FRG2C from chr 3 and FRG2C-1 from chr 22). Low consensus regions are shown in red.

**Supplemental Figure 2.**
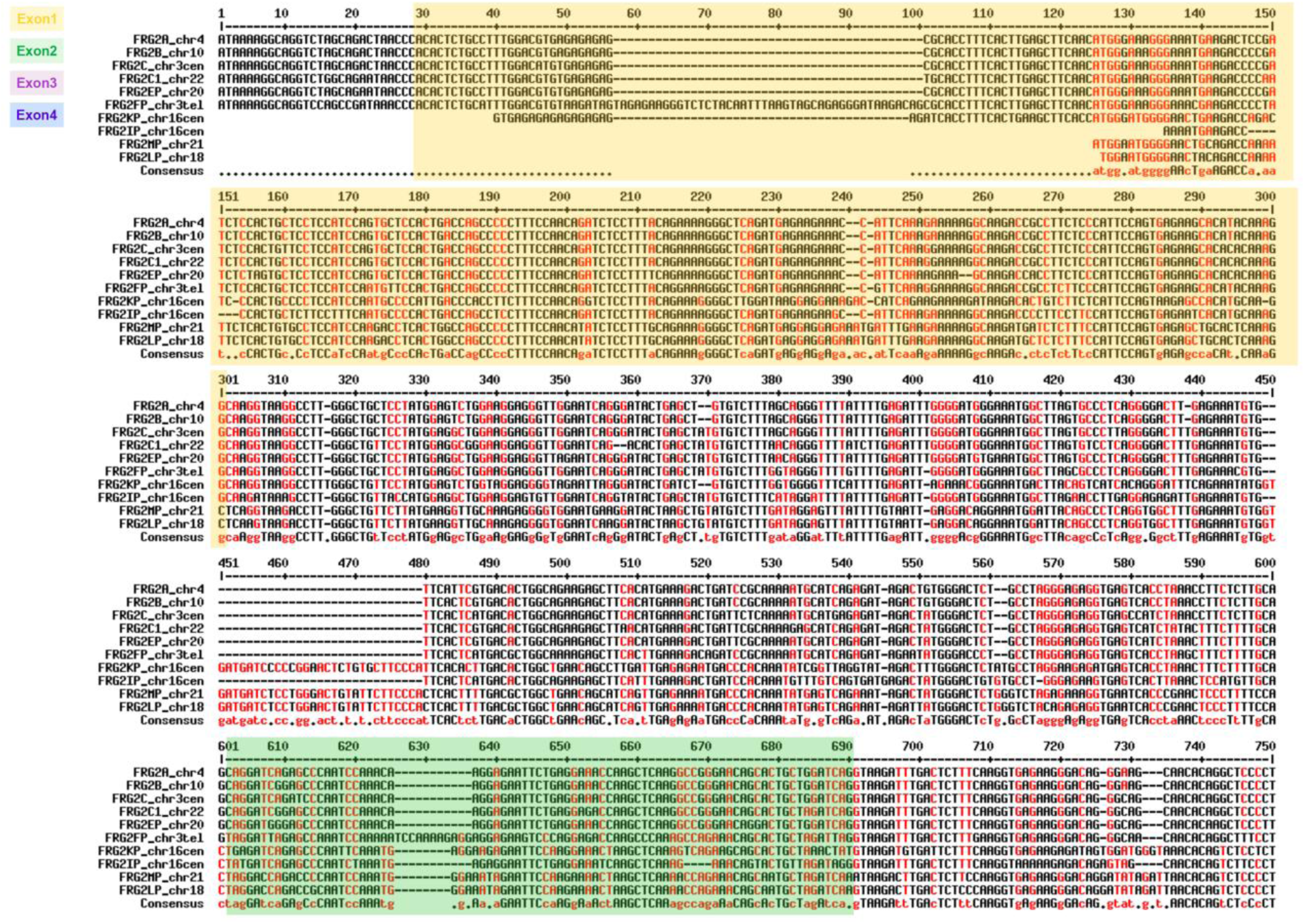
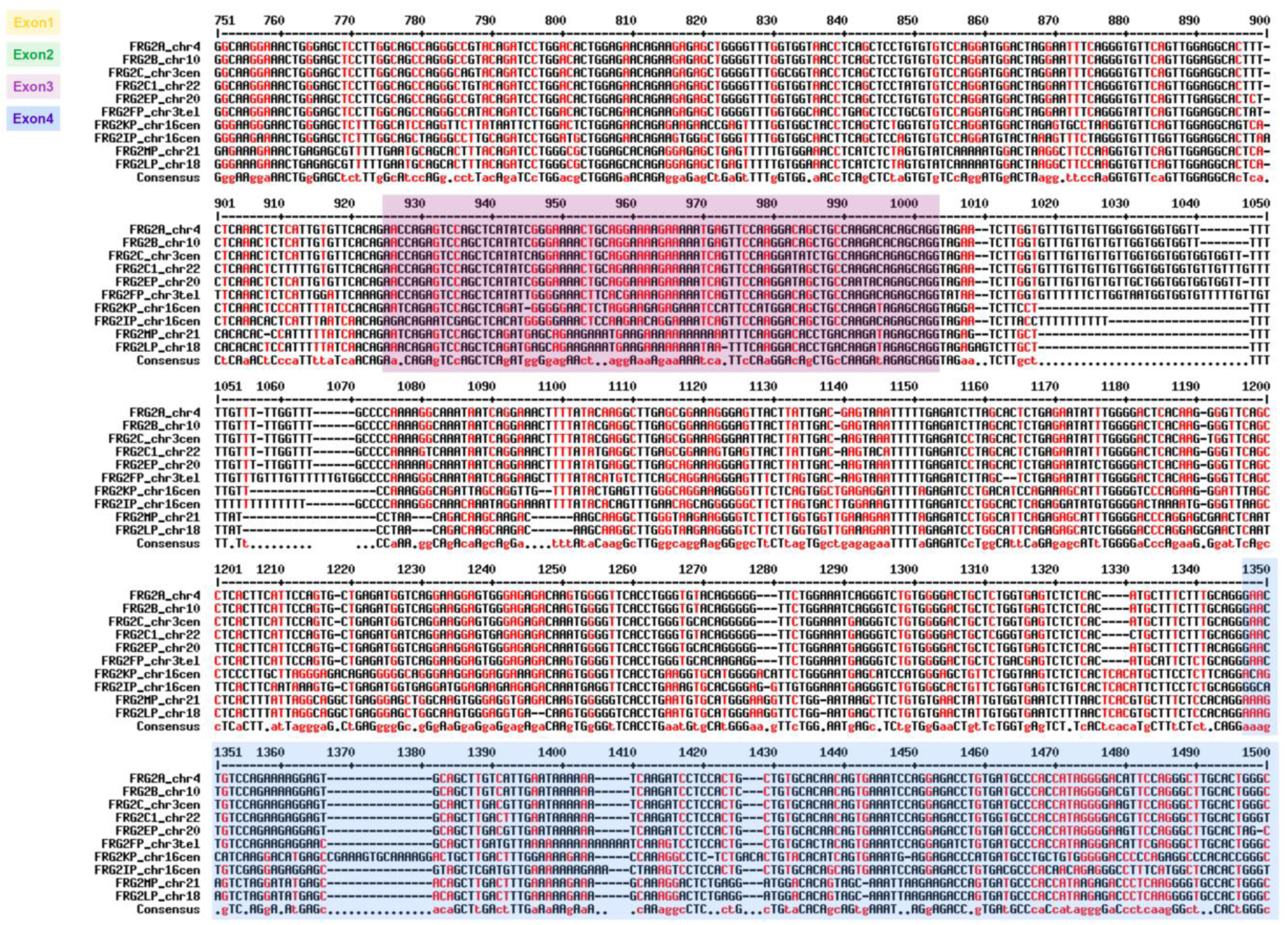
F*R*G2 paralogs sequences. Alignment of *FRG2* gene sequences of the complete copies from chromosomes 3, 4, 10, 20 and 22, and incomplete paralogs from chromosomes 16 (only two paralogs shown), 18 and 21. Low consensus regions are shown in red. The 4 exons of *FRG2A* on chr4 are shown as coloured boxes: yellow for exon 1, green for exon 2, purple for exon 3 and blue for exon 4.

**Supplemental Figure 3.**
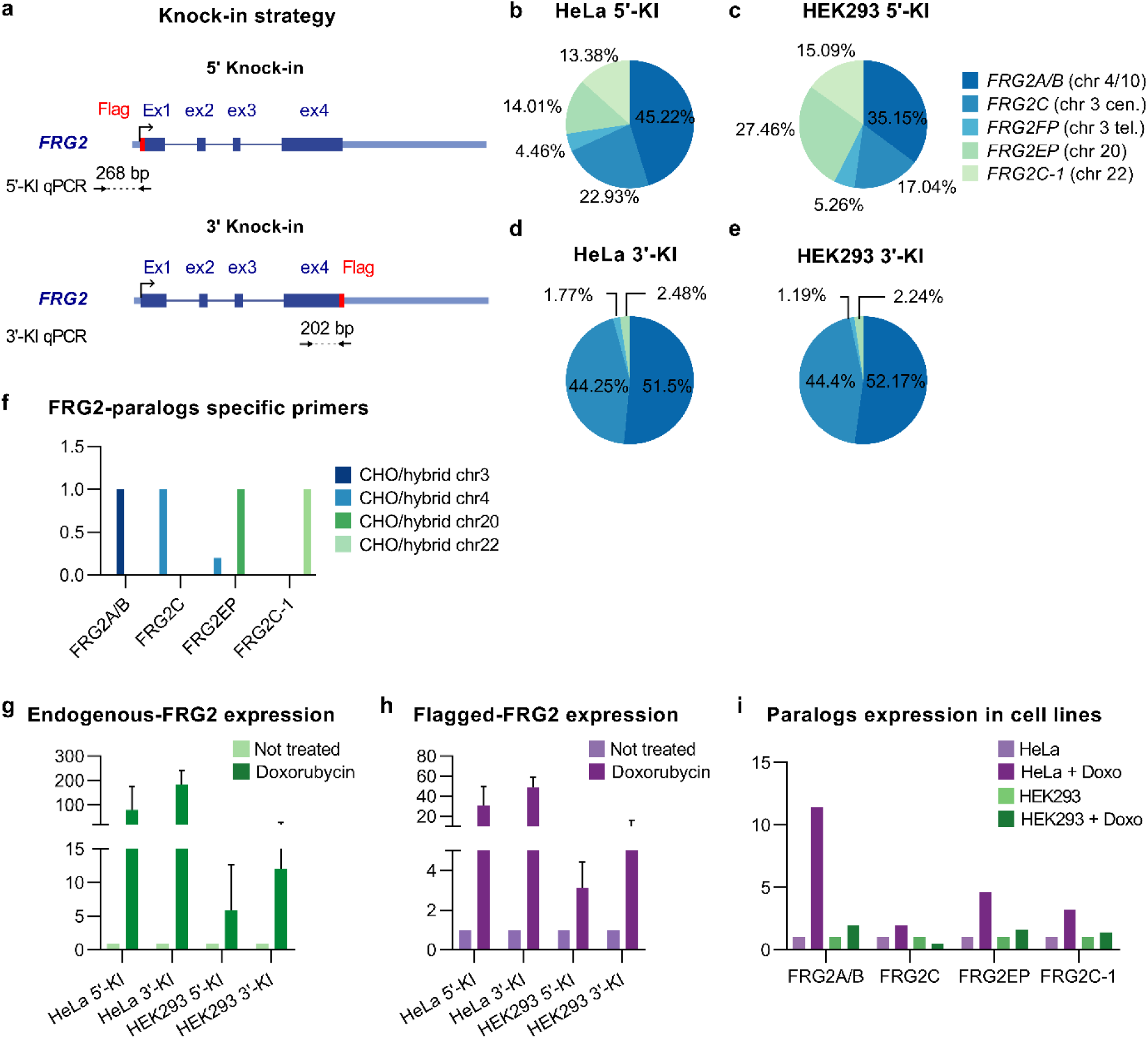
F*R*G2 knock-in strategy and editing validation. **a**, Schematic representation of the CRISPR-Cas9-mediated knock-in strategy designed to insert a Flag epitope at the N- and C-terminus of *FRG2* genes, both in HeLa and HEK293 cells. The two primer sets used to validate the efficacy of the editing are reported. **b-e**, Pie charts showing sequencing analysis performed after PCR-amplification of *FRG2* genes in CRISPR/Cas9 edited HeLa and HEK293 bulks. The analysis revealed that all the protein-coding genes (*FRG2A/B*, *FRG2C*, *FRG2C-1*) and also two non-coding copies from chromosomes 3 telomeric (*FRG2FP*) and 20 (*FRG2EP*) were well edited, in both cell lines. **f**, Validation by RT-qPCR of FRG2 paralogs-specific primers set. The experiment was performed by using cDNA obtained from CHO/hybrid of chromosomes 3, 4, 20 and 22, to validate the specific amplification of the different paralogs produced by the abovementioned chromosomes. **g**-**h**, RT-qPCR of cDNA obtained from HeLa and HEK293 edited bulks. The presence of the Flag-tagged FRG2s transcripts was detected by using a primer set able to amplify all the paralogs. cDNA was obtained from untreated and doxorubicin-treated bulks (1μg/mL, o.n. incubation). The expression of the endogenous FRG2s transcripts was assessed (g). The presence of the Flag-tagged FRG2s was confirmed and was similar to the expression of the endogenous (h), both at basal and overexpressed levels. **i**, RT-qPCR of FRG2 paralogs performed in wilde-type HeLa and HEK293 cells. Quantification of paralogs transcripts was evaluated also upon DNA damage induced by doxorubicin treatment.

**Supplemental Figure 4.**
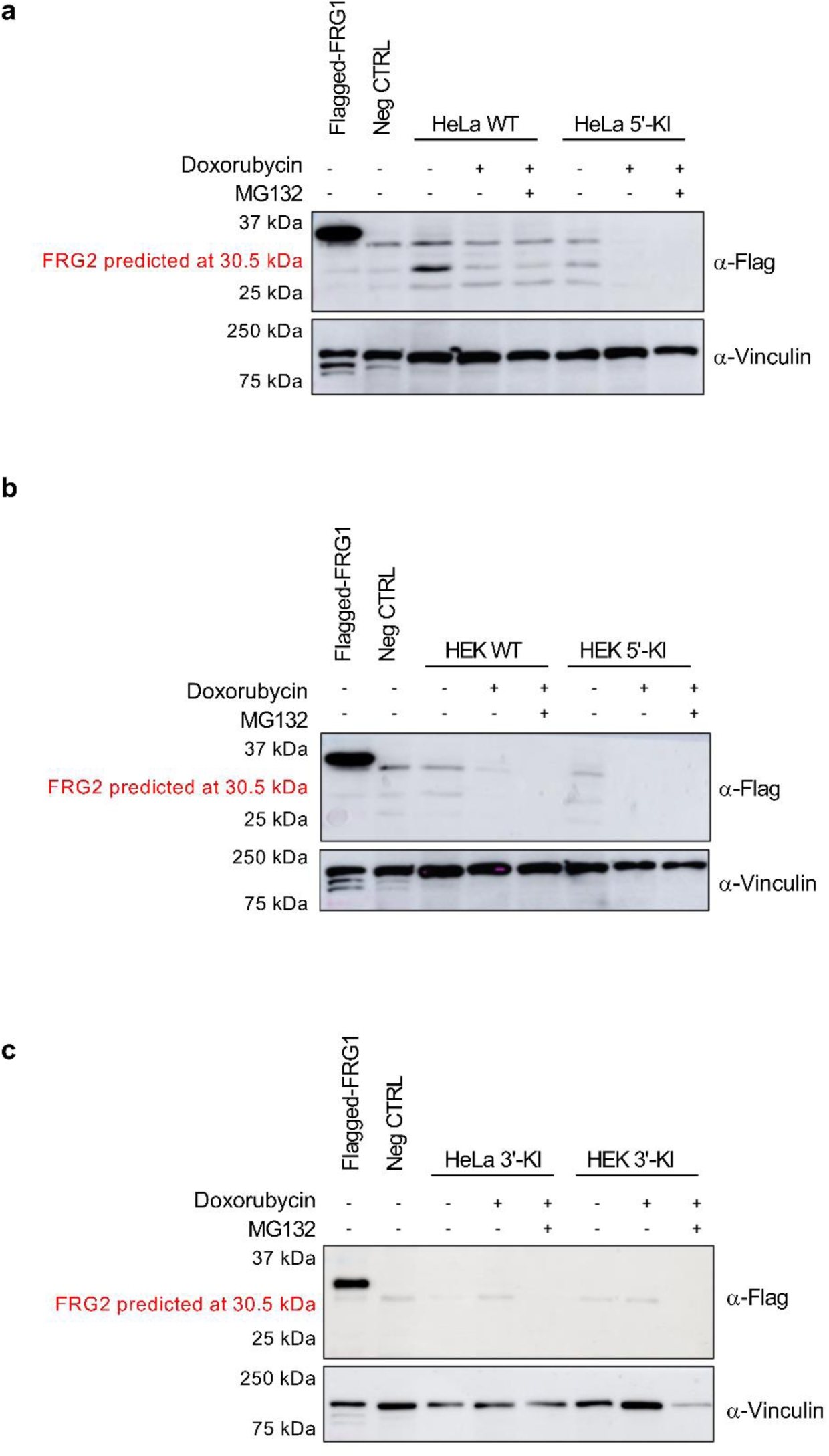
Western blotting against FLAG-FRG2 protein. **a-c**, Western blotting against FLAG-FRG2 fusion protein. Translation of the *FRG2* coding sequence is predicted to produce a 30.5 kDa protein. FLAG-tagged FRG1 protein extract was used as the positive control, while HeLa and HEK wild-type protein extracts (HeLa WT, HEK WT) were used as negative controls. Each bulk was assayed at basal condition, upon doxorubicin treatment alone and in combination with MG132^101^. Protein extracts from each bulk and experimental conditions showed equivalent results, with no detectable FRG2 protein.

**Supplemental Figure 5.**
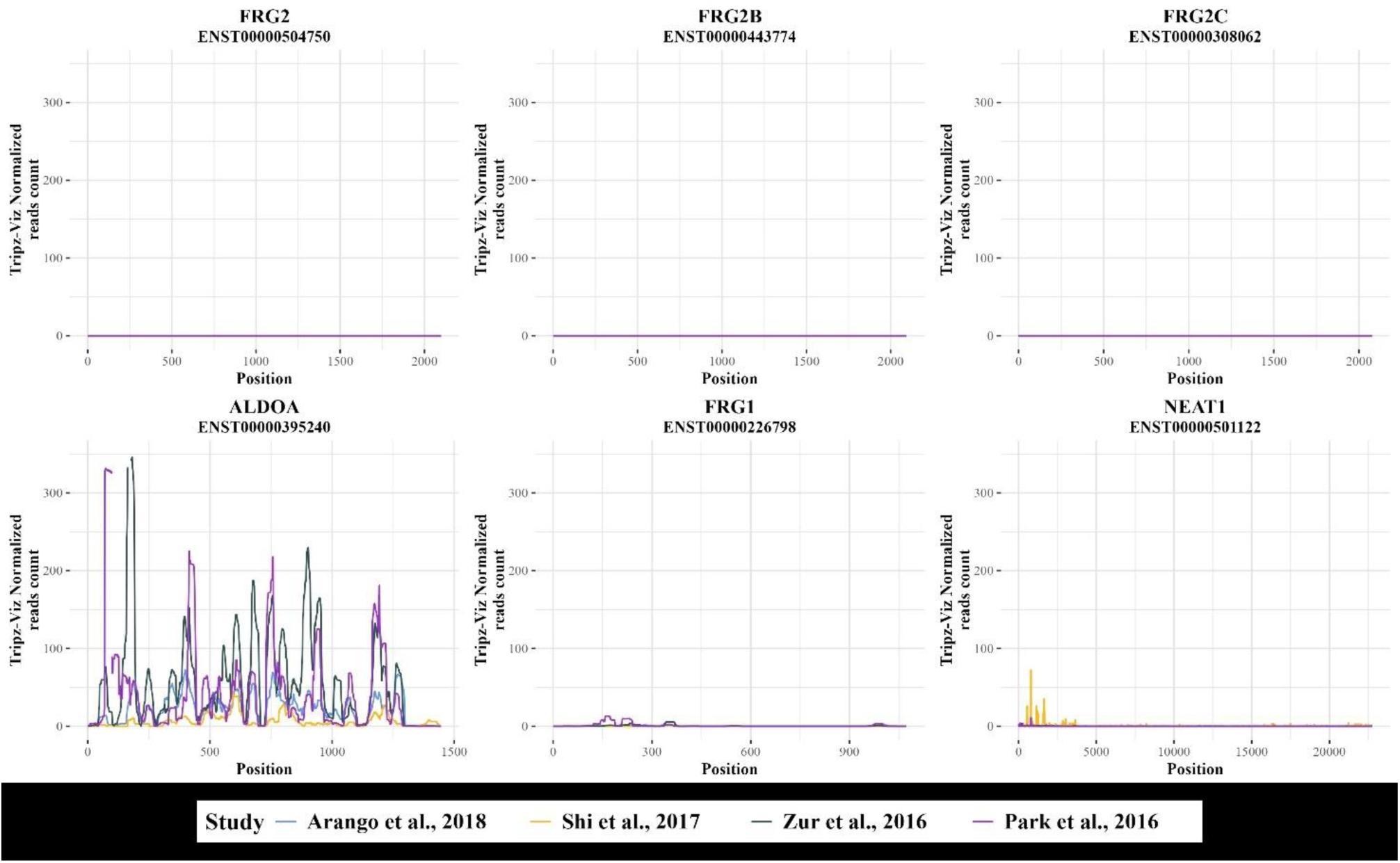
RiboSeq profiles of FRG2 paralogous genes, Aldolase (ALDOA), FRG1 and NEAT1. For each gene the tested transcript name is indicated as subtitle. Per each transcript the Ribosome footprint is shown as normalized read counts from the Tripz-Viz database. Lines are coloured according to the queried project^102–105^.

**Supplemental Figure 6.**
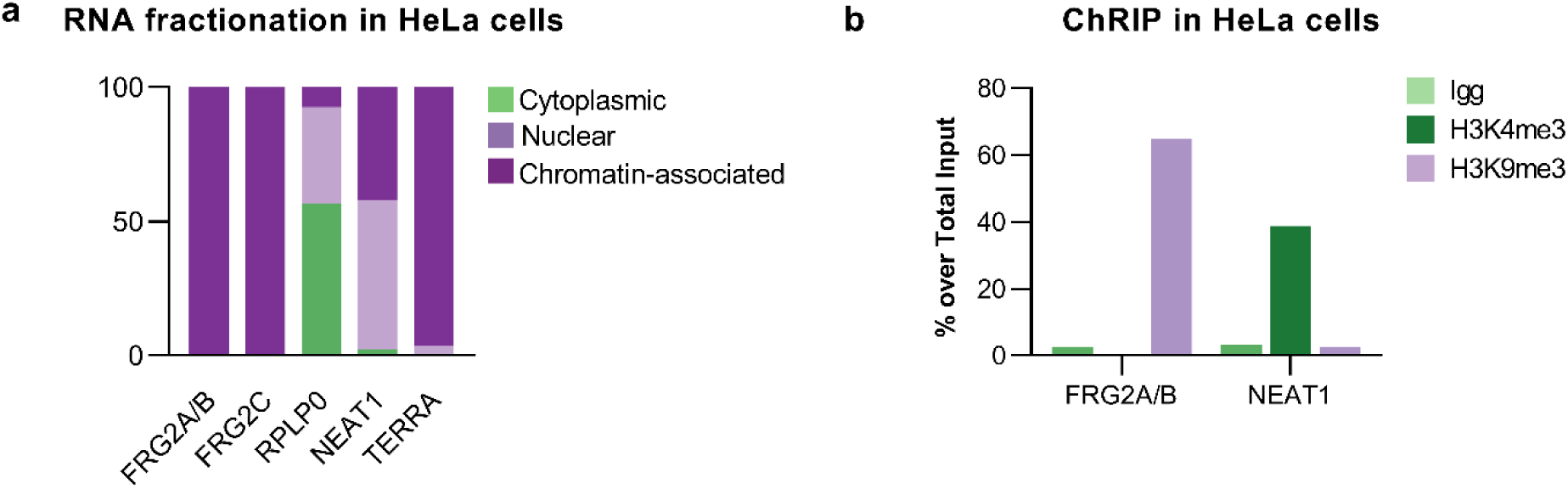
Analysis of FRG2 paralogs transcripts in HeLa cells. **a**, RT-qPCR of RNA fractionation experiment performed in HeLa cells. FRG2A and its paralogs transcripts were enriched in the chromatin associated fraction of RNA. RPLP0, NEAT1 and TERRA were used as positive controls to assess the enrichment of cytoplasmic, nuclear and chromatin-associated fractions, respectively^34,101^. **b**, Chromatin-RNA immunoprecipitation (ChRIP) performed in HeLa cells. FRG2A/B transcripts were enriched in the H3K9me3-immunoprecipitated materials. NEAT1 was used as a negative control since it is known to associate with H3K4me3-marked euchromatin^84^. Results are shown as percentages over total input.

**Supplemental Figure 7.**
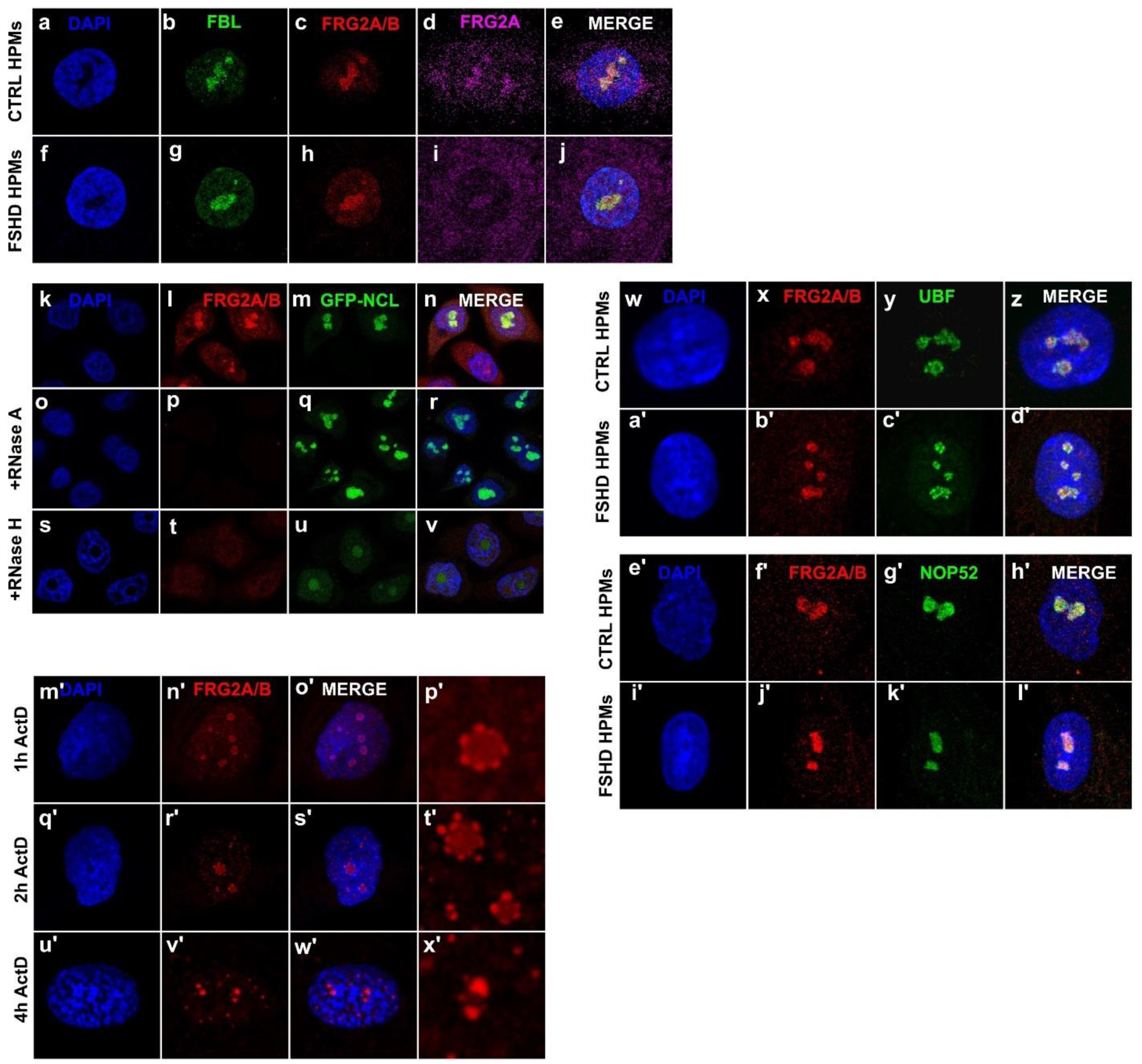
FRG2A transcript is localized to the nucleoli. **a-j**, immuno-RNA-FISH performed in CTRL and FSHD-derived human primary myoblasts (HPMs) with anti-FBL antibody (green), FRG2A/B specific fluorescent LNA probe (red) and FRG2A-only specific fluorescent LNA probe (magenta). The stain of the two different LNA probes is perfectly merged, confirming the localization of FRG2A within the nucleoli. **k-v**, RNA-FISH performed in HeLa Kyoto cells, a particular strain of HeLa stably expressing GFP-fused nucleolin (GFP-NCL, green). FRG2A/B specific fluorescent LNA probe (red) was used. Treatment with RNaseA (**o**-**r**) and RNaseH (**s**-**v**) reduced FRG2A/B fluorescent signal, confirming that FRG2A acts as an RNA molecule. **w-d’**, Immuno-RNA-FISH performed in HPMs obtained from FSHD patients and healthy relatives, with anti-UBF antibody (green) and FRG2A/B specific fluorescent LNA probe (red). Both in CTRL and FSHD cells, UBF and FRG2A/B are juxtaposed with no colocalization. **e’-l’**, Immuno-RNA-FISH performed in HPMs obtained from FSHD patients and CTRLs, by using anti-NOP52 antibody (green) and FRG2A/B specific fluorescent LNA probe (red). Both FRG2A/B and NOP52 signals are diffuse within the nucleolar body. **m’-x’**, RNA-FISH performed in HPMs treated with 1μg/mL of ACTD for 1, 2 and 4 hours. Upon increasing time of treatment, it is possible to appreciate the compaction of the nucleoli and FRG2A/B dots.

**Supplemental Figure 8.**
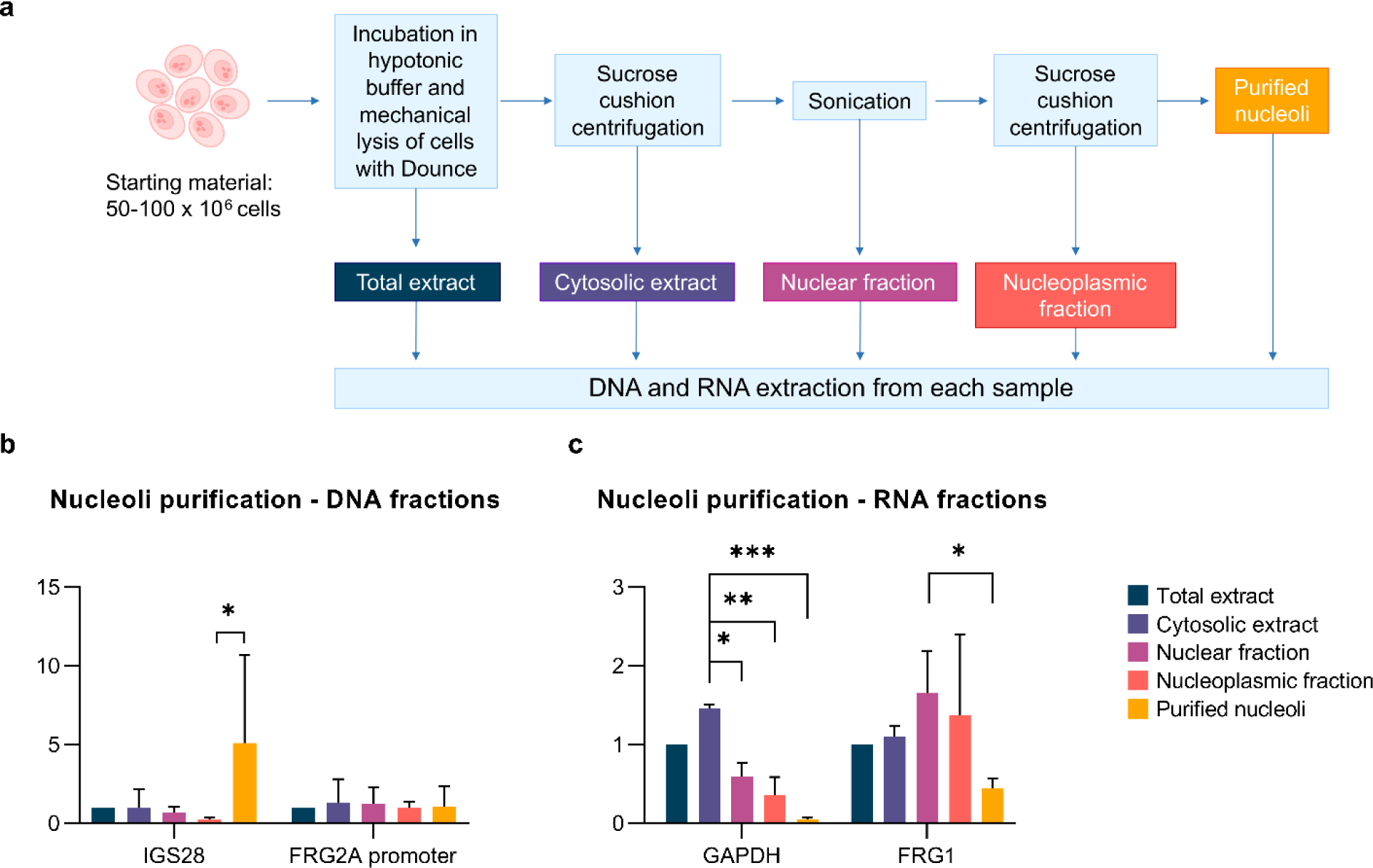
Nucleoli purification. **a**, Schematic diagram showing nucleoli purification protocol. **b**, RT-qPCR performed on DNA fractions obtained from the nucleoli purification protocol to verify the success of the assay. The enrichment of the specific nucleolar target (IGS28) in the purified nucleoli fraction was confirmed, while non-nucleolar region (*FRG2A* promoter) showed no enrichment in any of the fractions, as expected. Statistical significance was tested by using the two-way ANOVA analysis. p-value (****<0.0001). **c**, RT-qPCR performed on cDNAs obtained from the RNA fractions of the nucleoli purification protocol. As expected, protein-coding transcripts (GAPDH and FRG1) showed no enrichment in the nucleoli fraction. Statistical significance was tested by using the two-way ANOVA analysis. p-value (***<0.001).

**Supplemental Figure 9.**
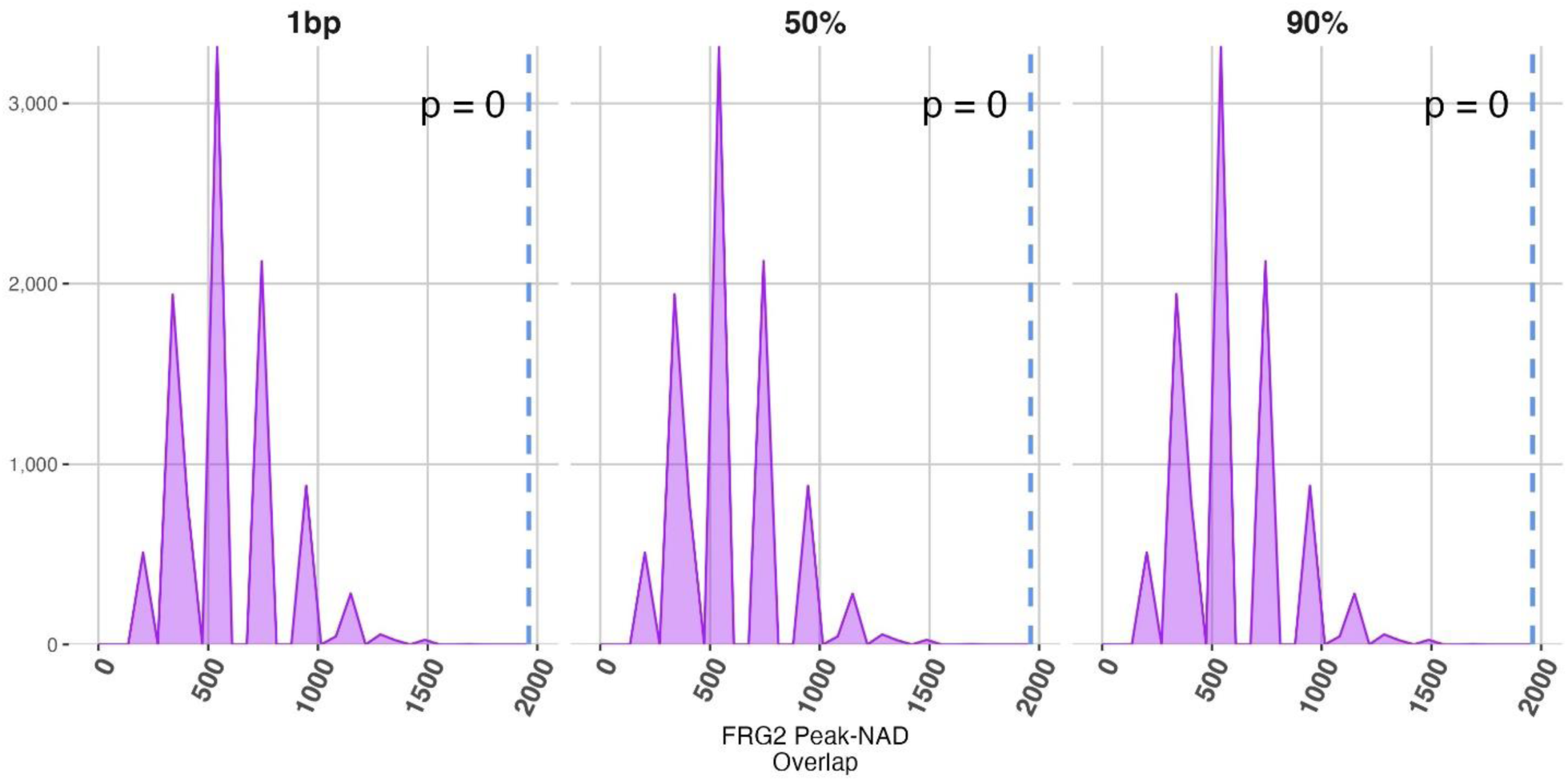
FRG2A/B association profile overlaps with NADs. By using bedtools random, 10,000 simulated bed files were generated using the same length distribution observed in inferred NADs. The co-localization of FRG2A/B peaks and simulated genomics intervals was measure by using three overlapping thresholds (1bp, 50% and 90% of FRG2 peaks intervals), and compared to the observed overlaps among FRG2A/B peaks and inferred NADs, by using Poisson distribution. The Density plot, shows the number of overlapping features and the counts on x and y axis, respectively. The dashed blue line corresponds to the overlaps measured among FRG2A/B peaks and inferred NADs. The p-value correspond to the provability to observe a greater number of overlaps in simulated data.

**Supplemental Figure 10.**
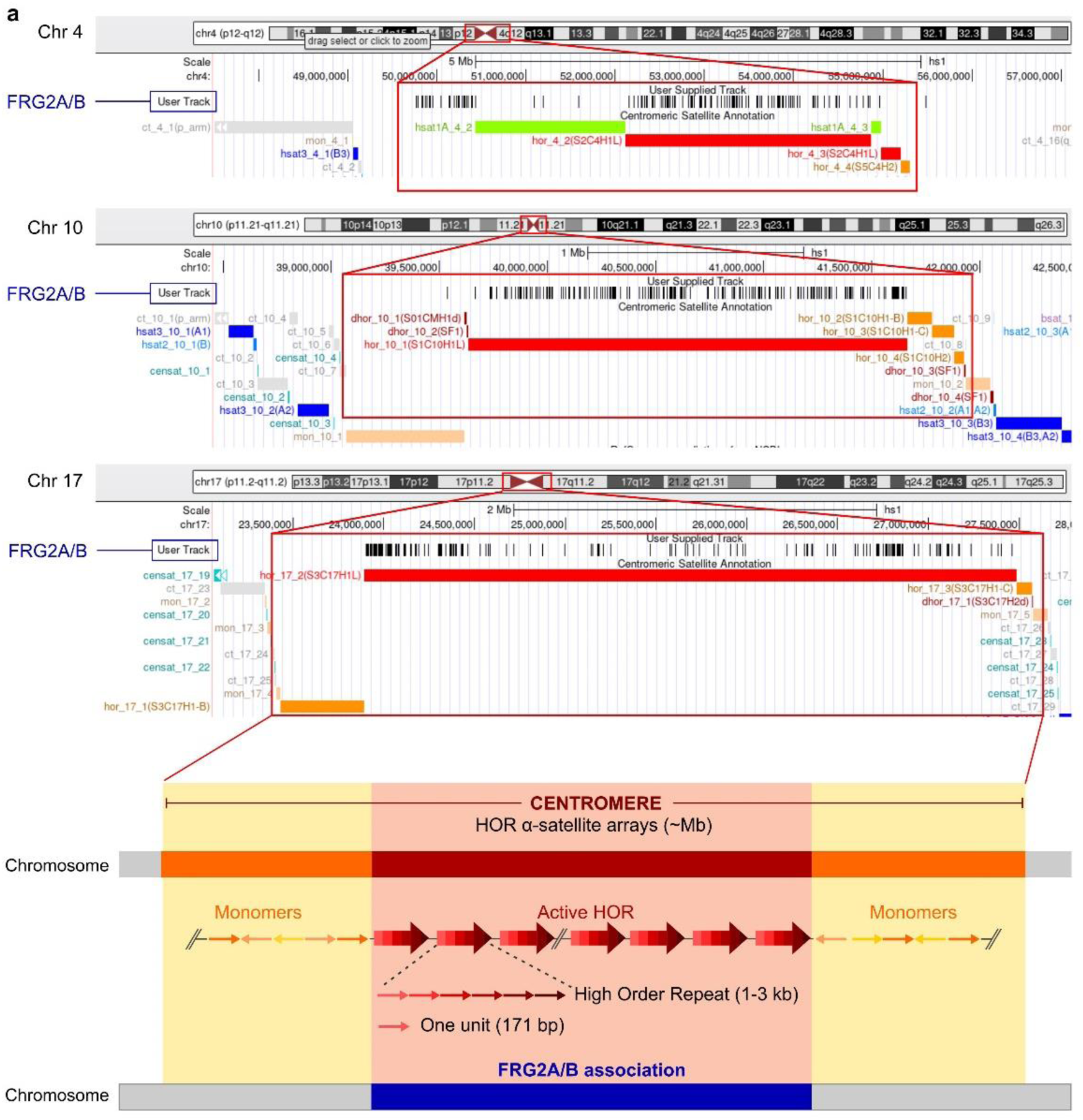
FRG2A/B associate with HORs. (Top panel) UCSC Genome Browser snapshot of human T2T centromeres from chromosomes 4, 10 and 17, which are the centromeres with the most FRG2A/B ChIRP reads. FRG2A/B ChIRP reads are shown in black. These reads fall within the red-boxed regions, which are enriched in Higher Order Repeat (HOR) regions and often flanked by satellite monomers (mon). (Bottom panel) Schematic representation of centromere composition, divided into monomers of α-satellites (orange) and α-satellites that comprise active HORs (red). FRG2A/B association is shown in blue, displaying its preferential association with the α-satellites of the active HORs, the site of centromere activity.

**Supplemental Figure 11.**
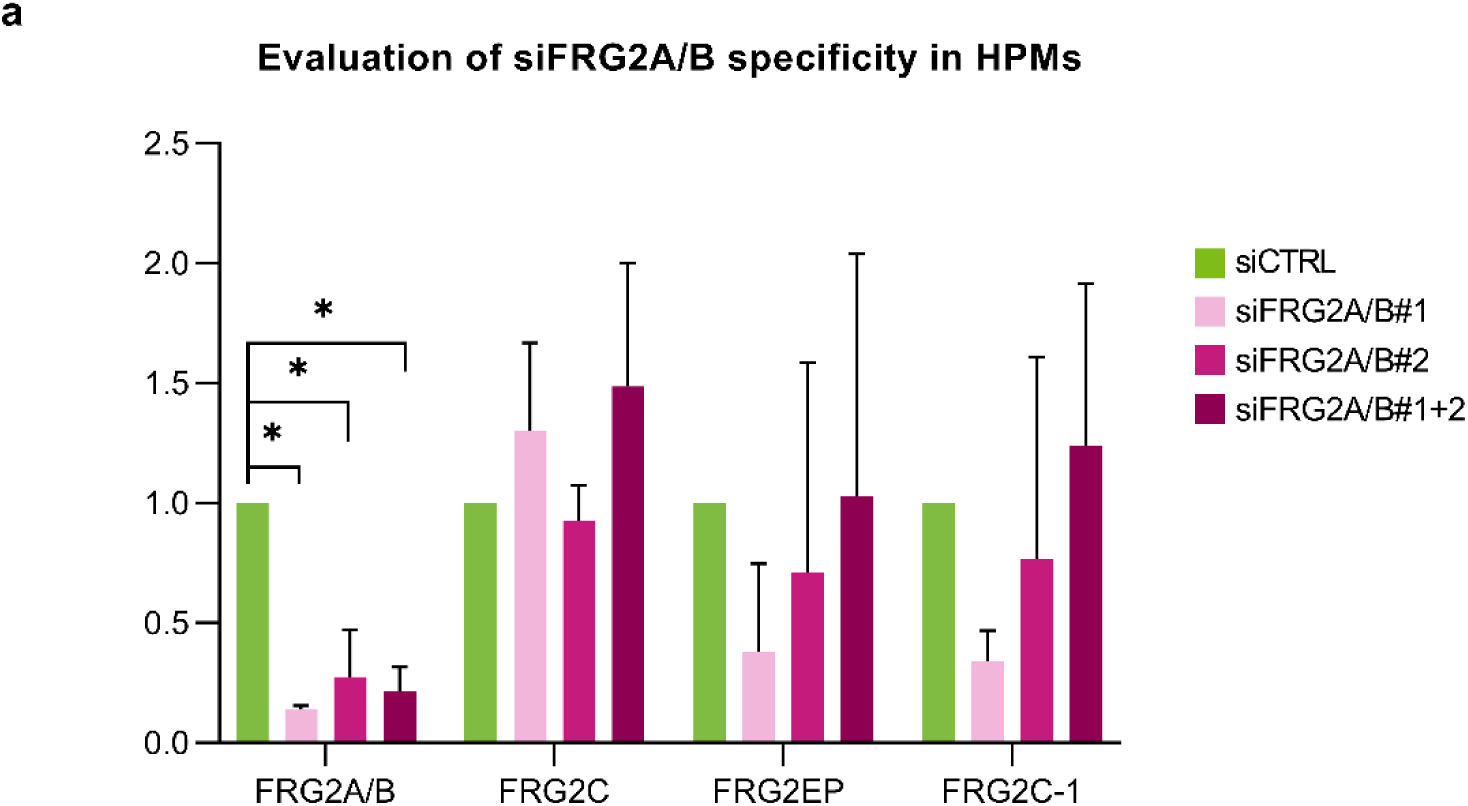
FRG2A/B gene silencing by short interfering RNA. RT-qPCR performed on cDNAs obtained from FSHD human primary myoblasts (HPMs) treated with siFRG2A/B#1 and #2, alone and in combination. Paralog-specific primer sets were used to evaluate the specificity of the siRNAs. Only FRG2A/B is significantly silenced after treatments. Statistical significance was tested by using the two-way ANOVA analysis.

**Supplemental Figure 12.**
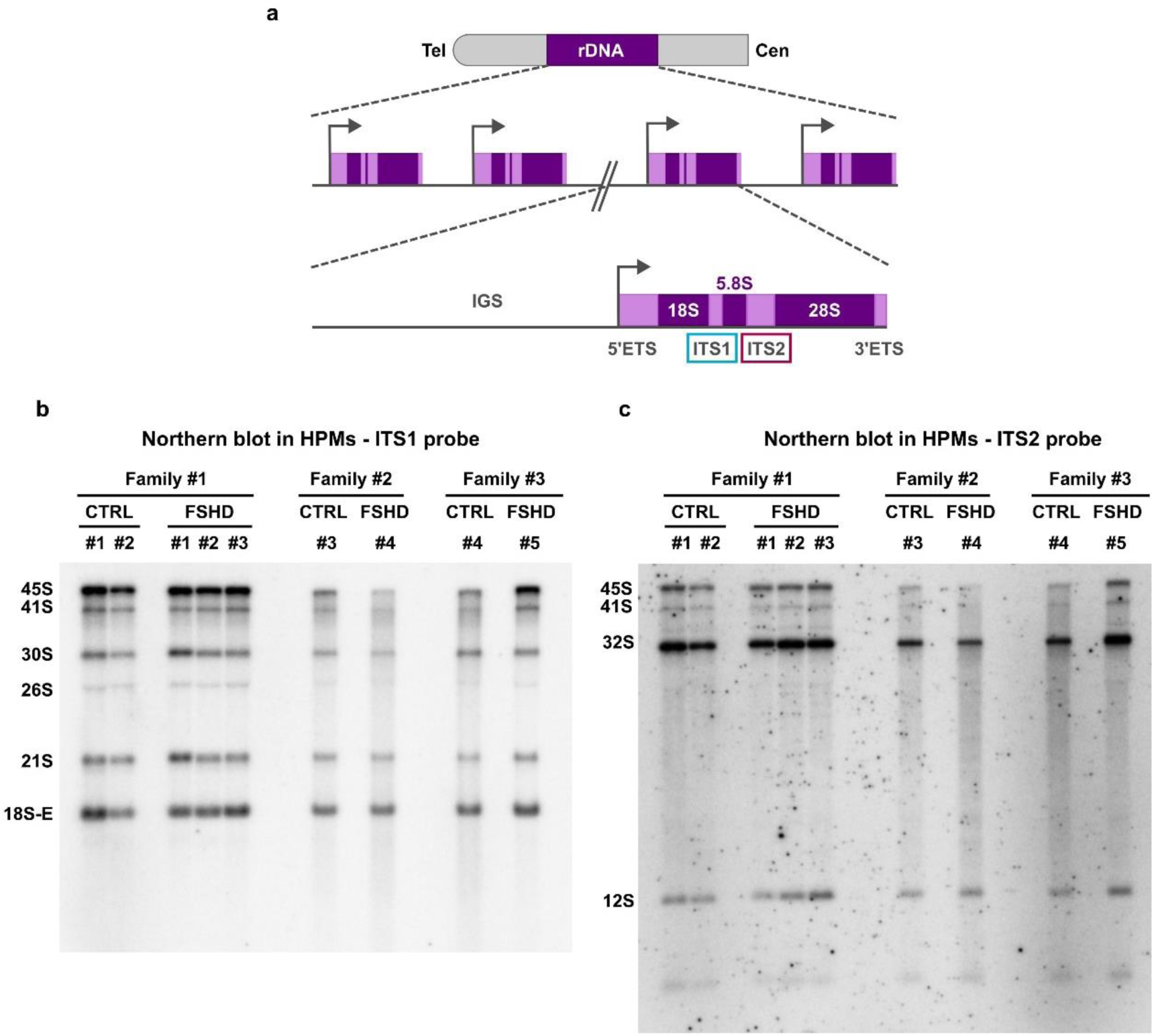
FRG2A/B does not affect rRNA processing pathways. **a**, Schematic representation of the rDNA arrays on the acrocentric chromosomes. Each rRNA Transcribed Region (purple) is spaced by the Intergenic Spacer (IGS) region. rRNA coding sequences are surrounded by the 5’-External Transcribed Spacer (5’-ETS) and 3’-External Transcribed Spacer (3’-ETS) and are internally spaced by Internal Transcribed Spacer 1 and 2 (ITS1 and ITS2, respectively). Position of probes used in northern blotting is indicated by blue and red squares corresponding to ITS1 and 2. **b-c**, Northern blotting assays of RNA extracted from CTRL and FSHD-derived HPMs. ITS1 and ITS2 probes were used to detect rRNA precursors of the 18S and 5.8S maturation pathways, respectively. Both assays did not reveal any defects in rRNA maturation processes in FSHD cells compared to controls.

**Supplemental Figure 13.**
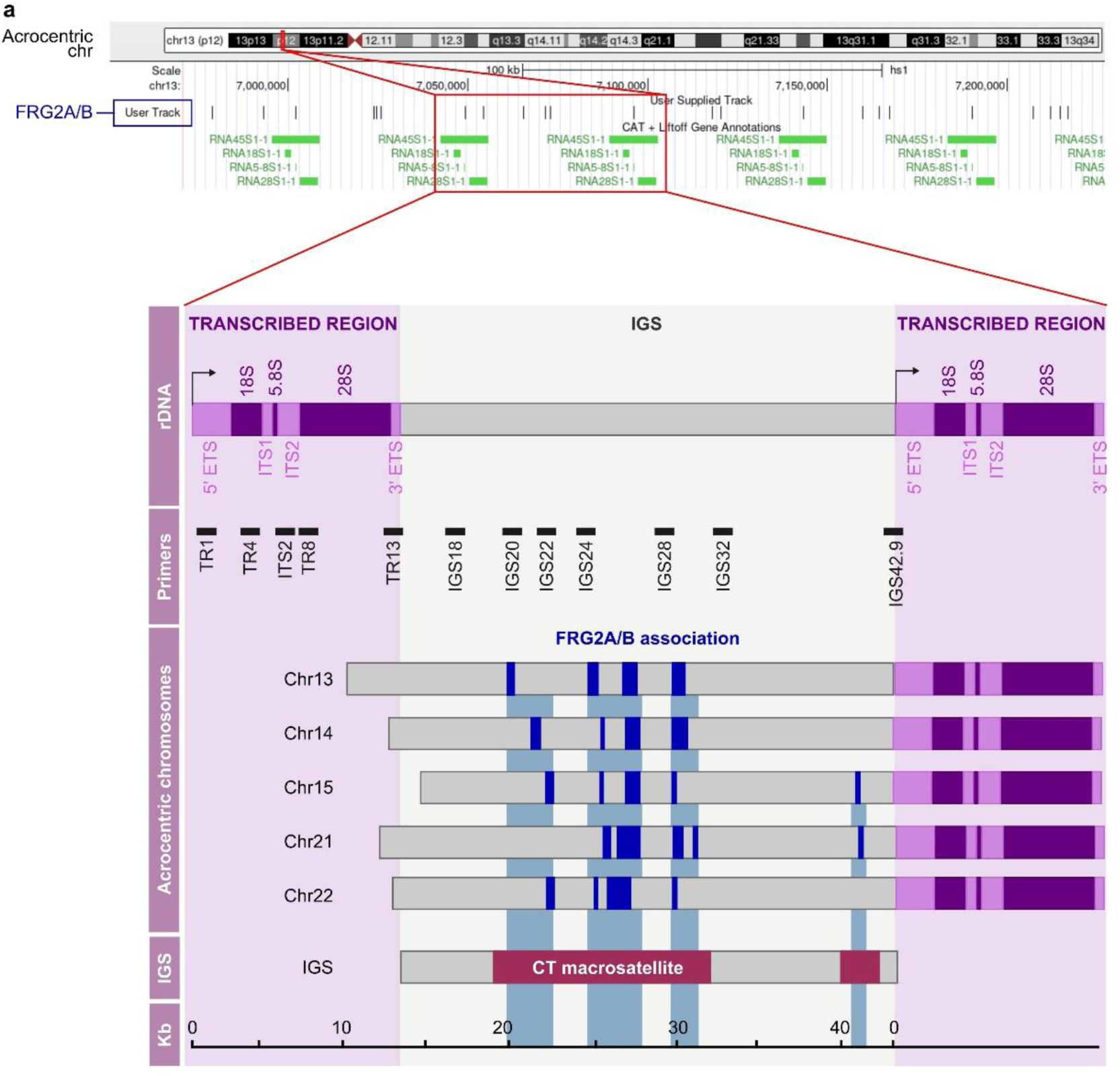
FRG2A/B-association with IGS at rDNA. UCSC Genome Browser snapshot of one of the acrocentric chromosomes, FRG2A/B track is reported in black as “user track”. Below is a schematic representation of the ribosomal DNA (rDNA) arrays, with two adjacent rRNA ORFs (purple) separated by the Intergenic Spacer (IGS) region (grey). The position of different primer sets used is reported, the names of which reflect the position of the amplified region within the 43 kb-long repeat. The five human acrocentric chromosomes are represented below, showing FRG2A/B-associated regions (in blue) as three distinct domains spanning from 20-22 kb to 30-32 kb. The CT-rich region within the IGS, named CT macrosatellite, is reported.

**Supplemental Figure 14.**
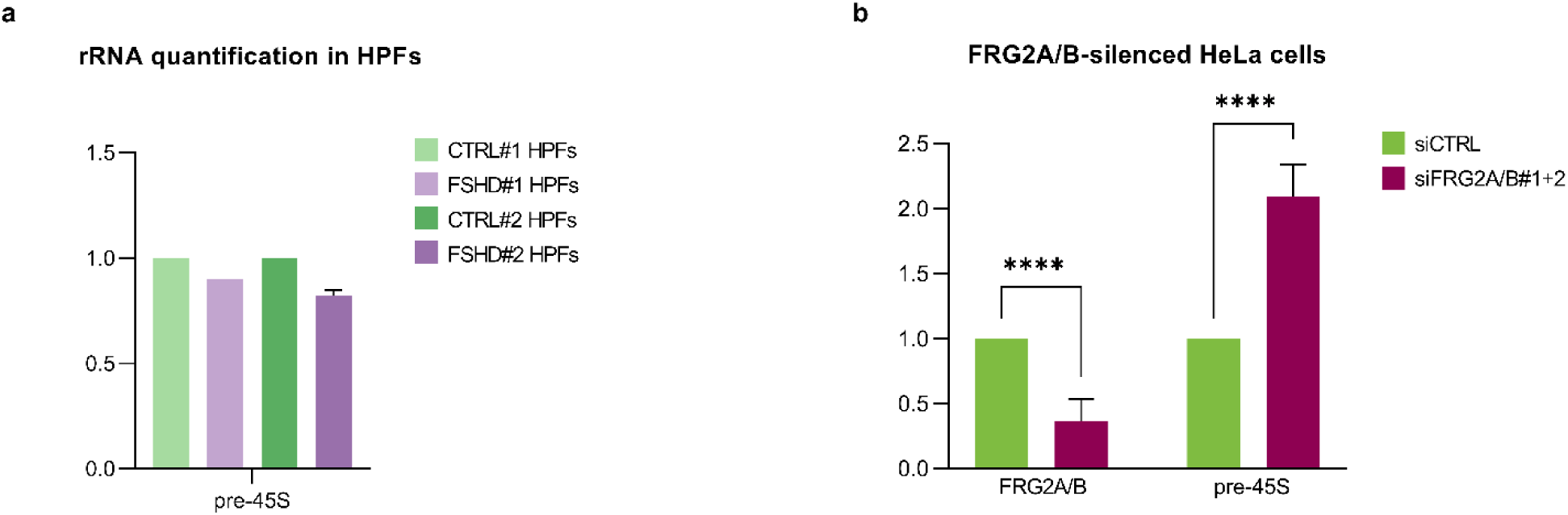
pre-45S transcription in HPFs and HeLa cells. **a**, RT-qPCR of pre-45S transcripts in human primary fibroblast (HPFs) shows no differences between FSHD patients and controls. **b**, RT-qPCR performed in siFRG2A/B-treated HeLa cells. Upon FRG2A/B silencing, pre-45S transcription is increased. Statistical significance was tested by using the two-way ANOVA analysis. p-value (****<0.0001).

**Supplemental Figure 15.**
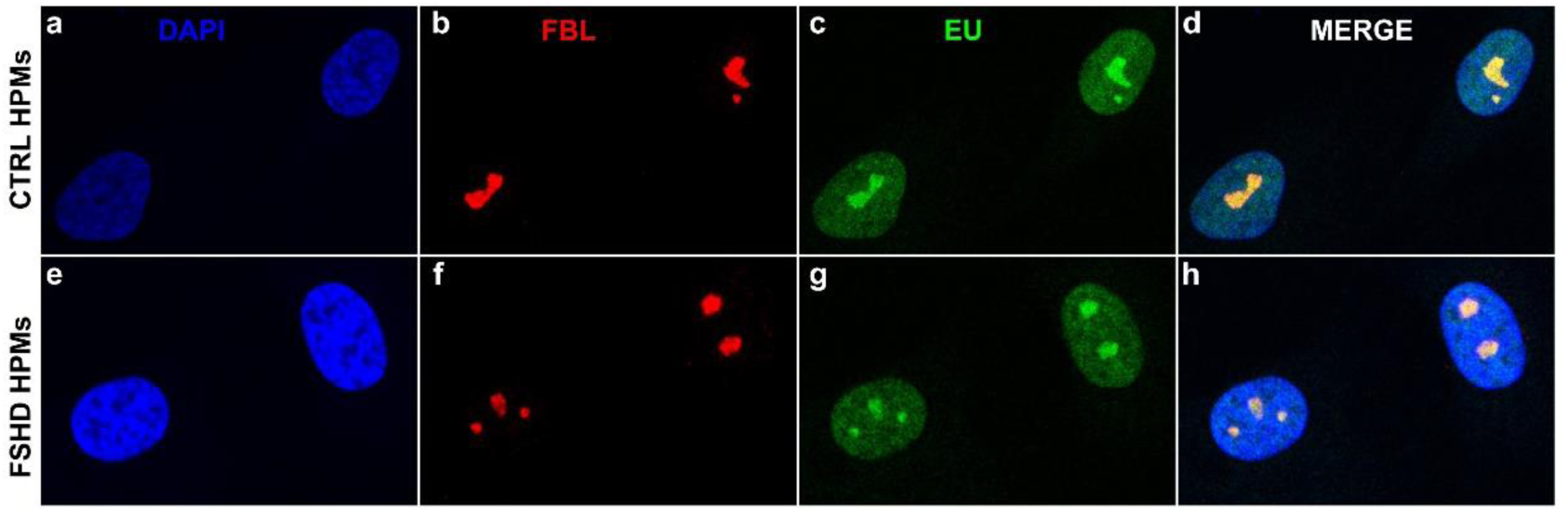
EU incorporation is reduced in FSHD HPMs. **a-h**, Immunofluorescence images showing 5-Ethynil Uridine (EU) incorporation assay performed in CTRL and FSHD in human primary myoblasts (HPMs). Incorporated EU was stained by a click-it chemistry (green), and nucleoli were marked by anti-FBL antibody (red).

**Supplemental Figure 16.**
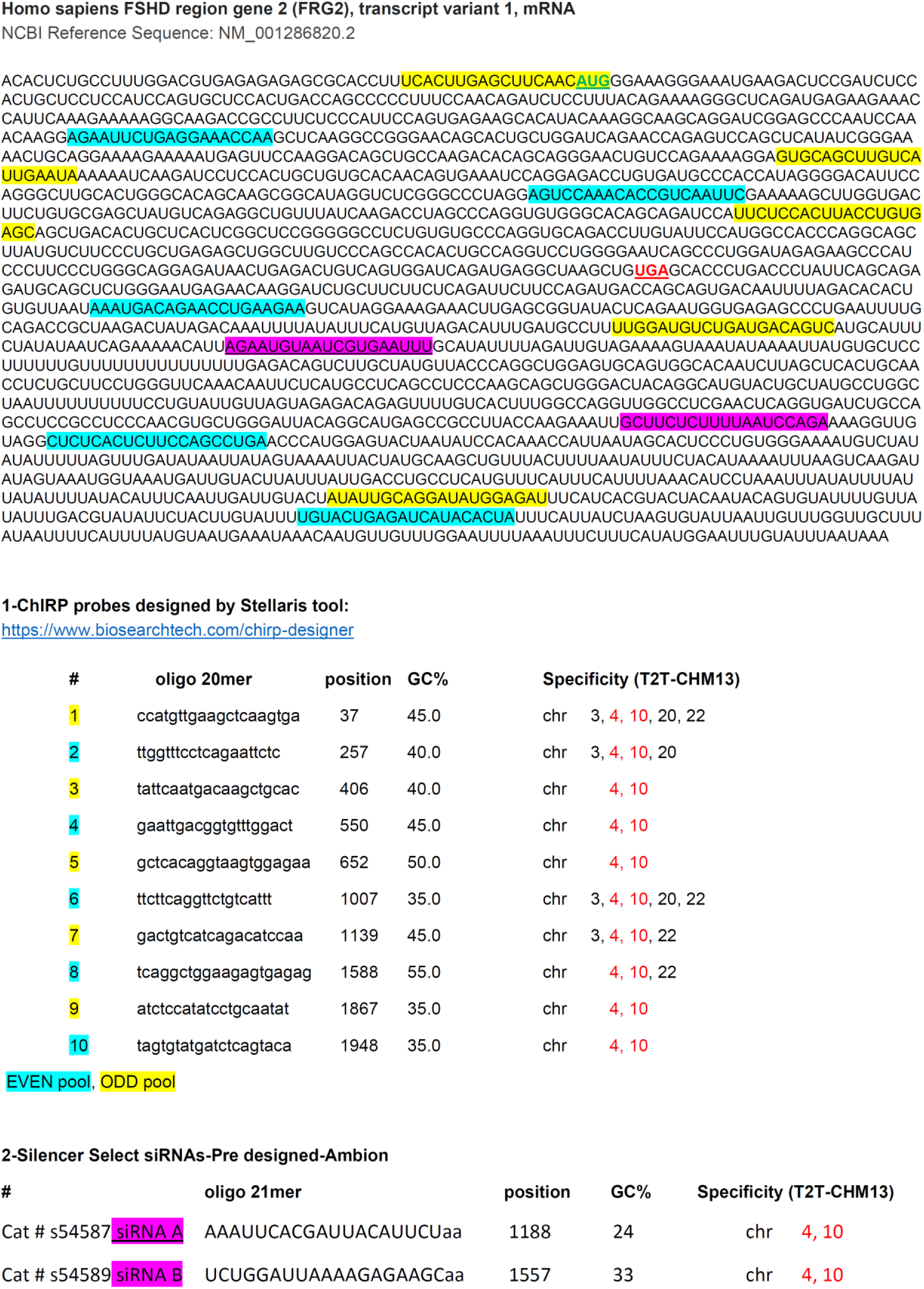
Homo sapiens FSHD region gene 2 (FRG2/FRG2A), transcript variant 1, mRNA, NCBI Reference Sequence: NM_001286820.2 and localization of specific probes used for ChIRP and silencing.

## SUPPLEMENTAL TABLES

**Supplemental Table 1. Clinical and molecular features of samples used in this study**: muscular biopsies, primary fibroblasts and primary myoblasts were obtained from FSHD families recruited through the Italian National Registry for FSHD (INFR).

**Supplemental Table 2. MS analysis in Hela cells**

**Supplemental Table 3. ChIRP data: WGS and RNA-seq**

**Supplemental Table 4. ChIRP-MS data**

**Supplemental Table 5. NADs annotation on T2T and FRG2 association profile overlap with NADs**

**Methods-Supplemental Table 1. List of primers and antibodies used in this study**

